# Thermodynamic limitations of metabolic strategies for PHB production from formate and fructose in *Cupriavidus necator*

**DOI:** 10.1101/2022.02.22.481442

**Authors:** Markus Janasch, Nick Crang, Manuel Bruch, Johannes Asplund-Samuelsson, Arvid Gynnå, Michael Jahn, Elton P. Hudson

**Affiliations:** Science for Life Laboratory, School of Engineering Science in Chemistry, Biotechnology and Health, KTH Royal Institute of Technology, P-Box 1031, 171 21 Solna, Sweden

**Keywords:** Cupriavidus necator, metabolic versatility, metabolic modeling, thermodynamics, PHB, Elementary flux modes, formatotrophy

## Abstract

The chemolithotroph *Cupriavidus necator* H16 is known as a natural producer of the bioplastic-polymer PHB, as well as for its metabolic versatility to utilize different substrates, including formate as the sole carbon and energy source. Depending on the entry point of the substrate, this versatility requires adjustment of the thermodynamic landscape to maintain sufficiently high driving forces for biological processes. Here we employed a model of the core metabolism of *C. necator* H16 to analyze the thermodynamic driving forces and PHB yields of different metabolic engineering strategies. For this, we enumerated elementary flux modes (EFMs) of the network and evaluated their PHB yields as well as thermodynamics via Max-min driving force (MDF) analysis and random sampling of driving forces. A heterologous ATP:citrate lyase reaction was predicted to increase driving force for producing acetyl-CoA. A heterologous phosphoketolase reaction was predicted to increase maximal PHB yields as well as driving forces. These enzymes were verified experimentally to enhance PHB titers between 60 and 300% in select conditions. The EFM analysis also revealed that metabolic strategies for PHB production from formate may be limited by low driving forces through citrate lyase and aconitase, as well as cofactor balancing, and identified reactions of the core metabolism associated with low and high PHB yield. The findings of this study aid in understanding metabolic adaptation. Furthemore, the outlined approach will be useful in designing metabolic engineering strategies in other non-model bacteria.

**Highlights:**

- Elementary flux modes of *C. necator* for PHB synthesis from fructose and formate.
- Metabolite sampling identified common reactions among EFMs with low driving force.
- PHB from formate shows low driving forces for aconitase, citrate lyase, NADPH synthesis.
- Phosphoketolase and ATP citrate lyase increased driving forces and PHB production.

## 1. Introduction

*Cupriavidus necator* H16 is an aerobic lithoautotrophic β-proteobacterium, known for its ability to produce large amounts of the bioplastic precursor polyhydroxybutyrate (PHB) (Ishizaki et al., 2001). While capable of metabolizing many compounds heterotrophically, *C. necator* can also utilize formate as the sole carbon and energy source. Formate is a promising substrate for biorefining, as it can be produced from CO_2_ via electrosynthesis (Claassens et al., 2019). In *C. necator,* formate is metabolized *via* formate dehydrogenases and the resulting CO_2_ is assimilated *via* the Calvin-Benson-Bassham (CBB) cycle (Friedrich et al., 1979). The combination of formatotrophic/autotrophic growth and a natural high flux toward acetyl-CoA-derived compounds such as PHB synthesis has renewed interest in *C. necator* as a potential autotrophic cell factory (Claassens et al., 2016; Krieg et al., 2018; Li et al., 2012; Panich et al., 2021). The significant metabolic flexibility of *C. necator* also makes it an ideal candidate to investigate metabolic adaptation. To utilize different substrates with different carbon and energy contents, microorganisms must adjust their metabolism, in particular the thermodynamic landscape that determines the directionality and driving forces of reactions, through changes in metabolite concentrations (Gerosa et al., 2015).

To elucidate the thermodynamics of metabolism, computational tools have been developed to integrate metabolite concentrations within metabolic network analysis (Asplund-Samuelsson et al., 2018; Gollub et al., 2021; Henry et al., 2007; Kümmel et al., 2006; Niebel et al., 2019). The addition of thermodynamic constraints in metabolic models can inform reaction directionality but also provide insight into complex metabolic phenomena. For example, reduced thermodynamic driving forces explain the reduced production capabilities of *Clostridia thermocellum* at high substrate loading due to accumulated hydrogen and formate (Thompson & Trinh, 2017). Oscillations in gas uptake rates during fermentation by *Clostridium autoethanogenum* were explained by oscillations in NADPH and thus oscillations in driving forces of key reactions (Mahamkali et al., 2020). Thermodynamics-constrained dynamic FBA was used to estimate internal reaction fluxes from extracellular metabolite profiles in flask and bioreactor cultivations of *C. necator* synthesizing PHB from glycerol (Sun et al., 2020). The comparison revealed that restricting TCA flux leads to increased PHB yields. In these cases, insights from thermodynamic analysis could suggest metabolic interventions to improve the limiting driving forces. One useful tool for analyzing pathway thermodynamics is the MDF (Max-min Driving Force) framework, an algorithm that maximizes the thermodynamically least favorable reactions in a pathway by optimizing metabolite concentrations (Noor et al., 2014) since reactions with low driving forces are expected to have high associated enzyme cost, and the directionality and flux through these steps are more likely to be controlled by substrate and product concentrations (Park et al., 2019).

Elementary flux mode (EFM) analysis computes minimal pathways throughout a metabolic network that can be analyzed thermodynamically (Gerstl et al., 2015; Hädicke et al., 2018; Jol et al., 2012). Any possible flux distribution can be retrieved by (non-negative) linear combination of elementary flux modes (EFMs) (Schuster et al., 1999), and flux distributions with maximum specific rate or optimized resource allocations have been found to be EFMs (Müller et al., 2014; Wortel et al., 2014). The combination of EFM analysis with thermodynamic evaluation via MDF is therefore a useful tool for analyzing metabolic networks such as that of *C. necator* where sparse fluxomics data (Alagesan, Minton, et al., 2018) means the *in vivo* flux distributions are not known or are variable. Thermodynamic analysis of EFMs explained production limits of *Clostridia thermocellum* at high ethanol titers by reversal or minimization of NADH generation by alcohol dehydrogenase (Dash et al., 2019). The MDF optimization algorithm however can provide non-unique solutions, so that in longer pathways, such as an EFM, there can be a degeneracy of metabolite concentrations that provide the same MDF value, or several reactions that operate at the calculated MDF. When performing MDF analysis for thousands of EFMs, this degeneracy can confuse the identification of what reaction is thermodynamically limiting.

In this study we used EFMs and thermodynamic analysis of a core *C. necator* network to explore routes to biomass and PHB from fructose and formate substrates. We enumerated EFMs connecting substrate and product, and investigated the thermodynamic driving forces within these using MDF analysis and random metabolite sampling. Additionally, we investigated the impact of two engineering strategies aimed at increasing the driving forces towards acetyl-CoA and thereby PHB production. The network was augmented with phosphoketolase reactions (XFPK1 and XFPK2) to source carbon from the Pentose Phosphate Pathway (PPP) and CBB cycle, and an ATP citrate lyase reaction (ACL), which is an ATP-driven reversal of the citrate synthase, and thus supplies acetyl-CoA from the TCA cycle (Kanao et al., 2002). By surveying all possible EFMs, we identified engineering targets which may increase PHB yield, as well as uncovered global “metabolic strategies” in *C. necator,* where certain reactions or pathways frequently constrain the thermodynamic landscape, regardless of the utilized substrate. Adding XFPK increased maximum attainable yields of PHB and driving forces towards acetyl-CoA on fructose and formate, while ACL only increased driving forces for EFMs to acetyl-CoA on fructose. Experimental testing of PHB yield from recombinant *C. necator* strains supported these predictions.

## 2. Model and methods

### 2.1 Metabolic network model

The metabolic network model was adapted from previous studies (“Experimental and Theoretical Analysis of Poly(β-Hydroxybutyrate) Formation and Consumption in Ralstonia Eutropha,” 2011; Lopar et al., 2014; Unrean et al., 2019), and modified with respect to the scope of the present study as described below. The final network covered the core central metabolism (Figure 1, 58 metabolic reactions). It includes the glycolytic Entner-Doudoroff pathway (ED), reactions for gluconeogenesis and the EMP pathway (note that *C. necator* H16 lacks a phosphofructokinase, PFK (Pohlmann et al., 2006)), TCA cycle as well as most reactions of the PPP (except 6-phosphogluconate dehydrogenase, 6PGD (Pohlmann et al., 2006)), which are largely shared with the CBB cycle. Note that re-capturing of CO_2_ using RUBISCO was allowed for fructose utilization, but there was no externally supplied CO_2_. The network also contained lumped reactions for the electron transport chain of oxidative phosphorylation, effectively capturing the obligate respiratory growth of *C. necator*. Based on the base model described above, an individual model variant was specified for both substrates (Figure 1A). Each model variant contained specific transport reactions for compounds taken up or excreted by the cell in addition to the core network. For the uptake of phosphate and formate, two alternative transport reactions were considered for each compound. Addition of exchange reactions for all metabolites potentially transported into and out of the cells allowed the steady-state balancing of these compounds beyond the system boundaries. For calculating maximum biomass and PHB yields and the production envelope *via* FBA for Figure 2, a single network model containing transport and exchange reactions for all substrates was used and constrained accordingly, see *2.4 FBA and production envelope*.

**Figure 1.**
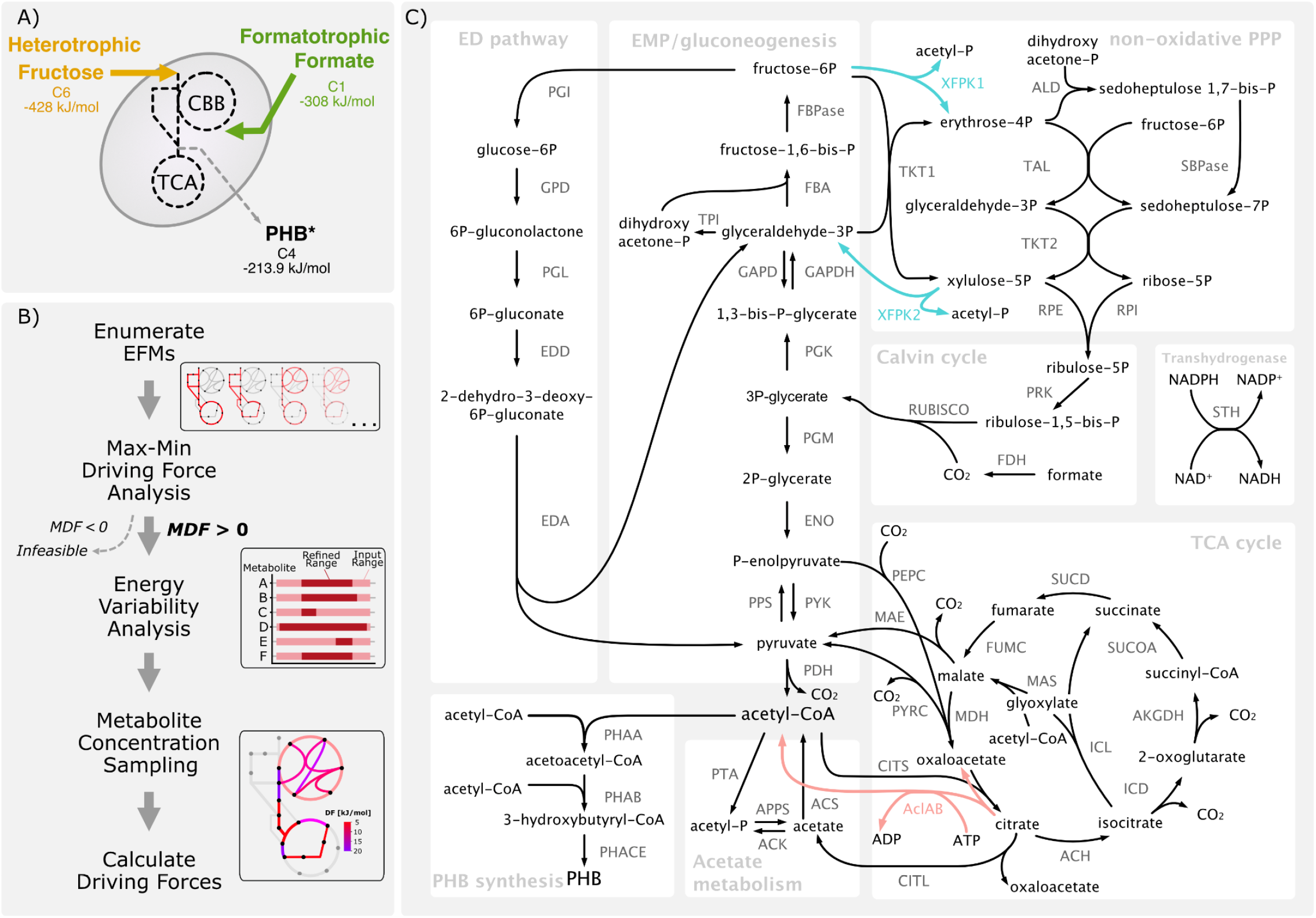
Analyzing the metabolic and thermodynamic landscape of *C. necator* and its dependence on substrate. **A)** The two different substrates investigated in this study with their number of carbon atoms, standard Gibbs energy of formation, and their entry point into metabolism. Each substrate imposes thermodynamic and stoichiometric constraints on potential biomass and PHB yield. *Properties of PHB are approximated from 3-hydroxybutanoate. **B)** Flow chart depicting analysis framework. **C)** Overview of metabolic network model used in this study. Heterologous reactions are marked in red and blue, respectively. Cofactors were omitted for better visibility except for the heterologous AclAB. Exchange and membrane-associated reactions are not shown. Arrows indicate default directionality used in the model and reaction names. Full reaction names are found in the supplementary information (Dataset S1).

**Figure 2:**
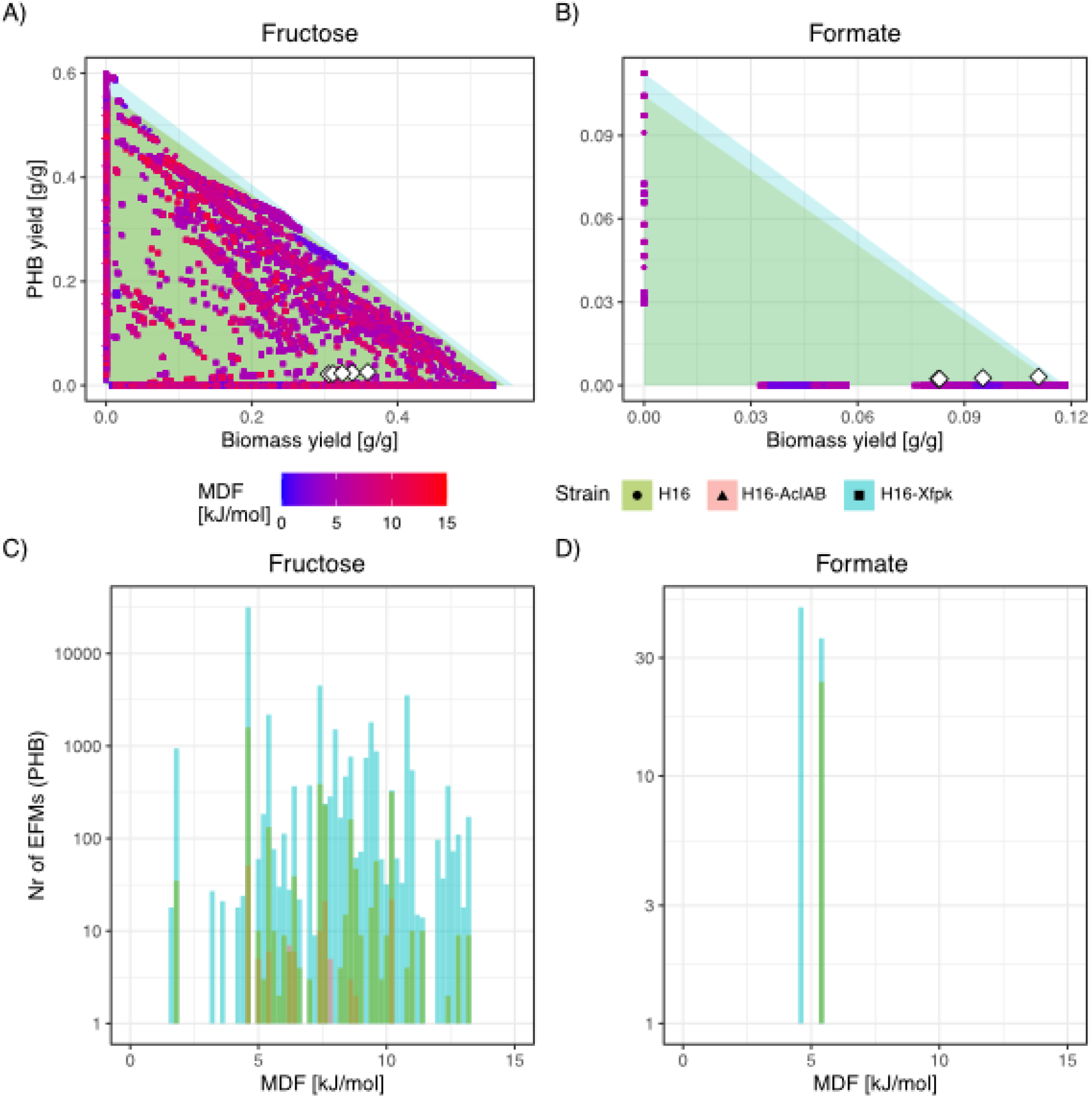
Overview of yields and MDF for EFMs reveal metabolic strategies. **A)** EFMs in the biomass and PHB yield space for the three strains on fructose. The shaded triangles correspond to the production envelopes calculated from FBA simulations. The color of each EFM corresponds to its MDF value. White diamonds indicate measured yields from chemostat experiments of the H16 wild-type at different dilution rates, taken from (Jahn et al., 2021). **B)** Same as A) but for formate. No EFMs using AclAB were found. **C)** Distributions of MDF values for each EFM producing PHB for the three simulated strains on fructose, revealing metabolic strategies with several EFMs having the same MDF. **D)** Same as C) but for formate.

### 2.2 Elementary Flux Modes and Thermodynamic Analysis

#### 2.2.1 Enumeration of Elementary Flux Modes

The thermodynamic landscape of metabolism depends in part on reaction directionality, which due to metabolic flexibility is difficult to know *a priori*. The uncertainty in reaction directionality can be circumvented by enumerating all possible flux distributions in the system by employing Elementary Flux Modes (EFMs). EFMs are non-reducible pathways, which form unique and linearly independent flux patterns, described in detail in (Zanghellini et al., 2013). Every possible flux distribution in a metabolic network can be constructed by linear combination of its EFMs. EFMs were enumerated using the EFMtool for MATLAB (Terzer & Stelling, 2008) [https://csb.ethz.ch/tools/software/efmtool.html].

#### 2.2.2 Max-Min Driving Force (MDF)

Lack of tight constraints in the EFM enumeration process can lead to combinations of reactions violating thermodynamic feasibility. To be thermodynamically feasible, a pathway or EFM has to consist exclusively of reactions with a positive Driving Force (DF), which corresponds to a negative change of Gibbs free energy, calculated via

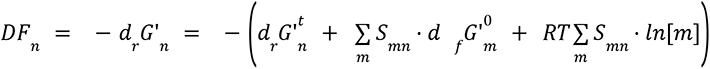

for reaction n, where *d_r_G’^t^_n_* is the Gibbs Free energy of metabolite transport across the cell membrane, *S* is the stoichiometric matrix, *d_f_G’^0^_m_* is the standard Gibbs Free energy of formation of compound m, *R* is the universal gas constant, *T* is the temperature and *ln*[*m*] is the natural logarithm of the concentration of compound m.

To ensure that an EFM consists only of reactions with a positive DF, the Max-Min Driving Force (MDF) algorithm was employed (Noor et al., 2014). The MDF framework is a linear optimization algorithm that, given directionality and *d_r_G’^0^* of reactions and concentration bounds of metabolites, finds a set of metabolite concentrations so that the reaction with the lowest DF in a pathway, marking a potential thermodynamic bottleneck, is as far from equilibrium as possible. The linear programming problem is formulated as:

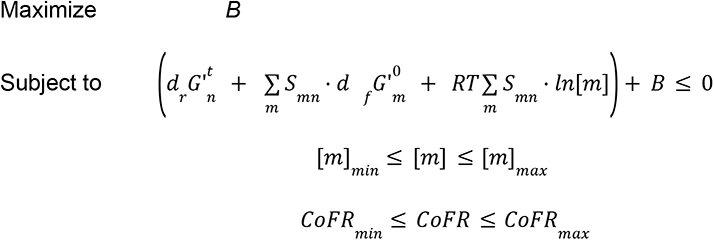

with *CoFR* being the ratio of the cofactor pairs NADH/NAD, NADPH/NADP, ATP/ADP and quinol/quinone, and *B* being the resulting maximum minimum driving force (MDF) value of a pathway. Several reactions in a pathway can operate at the MDF. Gibbs Free energies of formation *d_f_G’^0^*_m_ were retrieved from eQuilibrator 2.2 (Flamholz et al., 2012), with an internal pH of 7.5, ionic strength of 0.25, Gibbs Free energy of metabolite transport were calculated according to (Jol et al., 2010), see Supplementary Methods 1. Internal metabolite concentration ranges were set between [*m*]*_min_* = 1 µM and [*m*]*_max_* = 10mM, with the exception of glutamate (glut_c [*m*]*_max_* = 100 mM) (Bennett et al., 2009). Details about the formulation of the linear programming problem are found in Supplementary Methods 2. A negative MDF renders an EFM thermodynamically infeasible. Biomass formation and the final step of PHB synthesis were assumed to be sufficiently thermodynamically forward driven under all conditions and were excluded from the MDF analysis. The driving force of biomass formation is challenging to determine and can differ vastly, even in more detailed metabolic models (Saadat et al., 2020).

#### 2.2.3 Energy and concentration variability analysis

While the resulting MDF value marks the unique optimum value for each EFM, there can exist many metabolite concentration sets that lead to this MDF. To explore the potential variability in metabolite concentrations leading to the same MDF value, metabolite concentrations were minimized and maximized within an MDF window of 0.9x to 1.0x of the optimum to account for uncertainties associated with standard Gibbs energies of formation for the compounds.

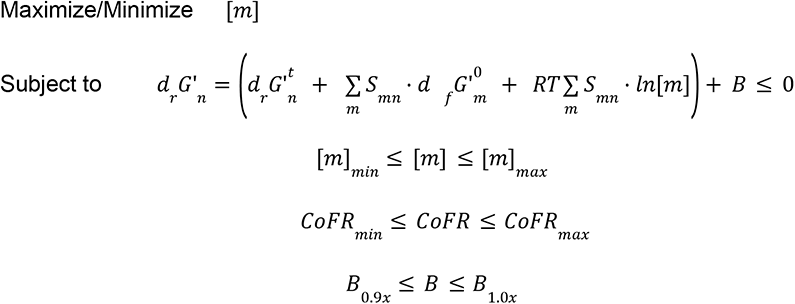

with *B_1.0x_* being the previously determined optimal MDF solution (2.2.2) and *B_0.9x_* being 90% of the optimal solution.

Similarly, the change of Gibbs free energy of each reaction can be evaluated:

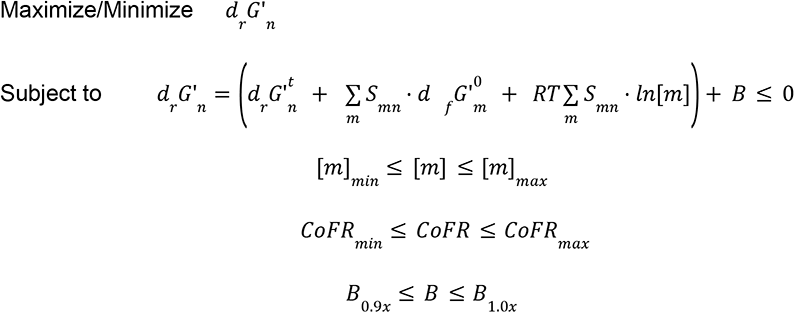

#### 2.3.4 Computational sampling of metabolite concentrations for driving force estimation

The metabolite concentration ranges of the variability analysis above were used to constrain the solution space for a hit-and-run style random sampling of metabolite concentrations to get a probabilistic overview of DFs for each reaction in each EFM, constrained by the MDF. The hit-and-run sampling starts at the metabolite concentrations determined by the MDF analysis, picks a random direction in which to modify the vector of metabolite concentrations, and determines to what amplitude the modification may be performed in the positive and negative directions without violating the concentration and driving force constraints. A random amplitude of change within the determined limits is added to the previous set of metabolite concentrations to perform one step of hit-and-run sampling. For each EFM, 5000 metabolite concentration sets were created through the stepwise hit-and-run sampling and subsequently the DFs of each reaction were calculated. For comparison between EFMs, the median DF and the median absolute deviation (MAD) from the DF for each reaction was calculated. The hit-and-run algorithm is supplied as a Python script at https://github.com/MJanasch/ThermoCup.

### 2.3 Yield and MDF scores

A reaction scoring system was devised to determine the influence of individual reactions on yields and MDF values. The reaction score compared the mean biomass and PHB yield as well as the MDF of the EFMs using a certain reaction with the means of these values over all EFMs. Reactions more often used within EFMs with higher yields or MDF, respectively, will have a positive score and vice versa.

Scores were calculated according to the formula:

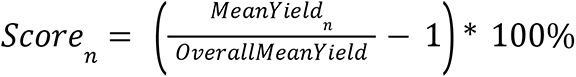

Where the score of reaction n is a measure for the ratio between the mean yield (or MDF) of all EFMs using reaction n (*MeanYield_n_*) and the mean yield (or MDF) of all EFMs (*OverallMeanYield*).

### 2.4 FBA and production envelope

Maximum biomass and PHB yields were calculated for each condition, with exchange reactions for the corresponding substrate fixed at 1 mmol⋅gDW^-1^⋅h^-1^. First, the objective function was set to maximizing biomass production. Second, the obtained optimal value was set as a fixed constraint (lower and upper bound) and successively multiplied with values linearly decreasing from 1 to 0, while the objective function was set to maximize PHB production. Obtained values for both biomass and PHB production were normalized with the molecular weight of the substrate. This process was repeated for each of the substrates, and the resulting pairs of biomass and PHB form the production envelope in Figure 2A and B.

### 2.5 Construction of *C. necator* mutant strains

Three *Cupriavidus necator* mutant strains were developed and utilized in this work, expressing either red fluorescent protein (Rfp), the ATP citrate lyase from *Chlorobium limicola* (AclAB, Uniprot Q9AJC4 and Q9AQH6), or the X5P/F6P phosphoketolase from *Bifidobacterium breve* (Xfpk, Uniprot D6PAH1). Transgenes were expressed from an episomal vector under the control of the arabinose-inducible pBAD promoter (Alagesan, Hanko, et al., 2018). The episomal vector was pBBR1MCS-2, a gift from Kenneth Peterson (Addgene plasmid #85168; https://www.addgene.org/85168/; RRID:Addgene_85168).

The AraC activator and P_BAD_ promoter were synthesized as a single dsDNA fragment by Integrated DNA Technologies, following the sequence reported by Alagesan (Alagesan, Hanko, et al., 2018). All three transgenes were codon optimized for expression in *E. coli,* and synthesized as either dsDNA fragments or gene clones by Integrated DNA Technologies. The Xfpk had a 6-His tag on the N terminus. To assemble the vectors, Hi-Fi assembly was used. Gene maps, vector sequences, strain and primer lists can be found in Supplemental Material (Figure S11, Table S3-5). The arabinose inducible vectors expressing Rfp, aclAB and Xfpk were termed HSLP 001, HSLP 002 and HSLP 003. The *C. necator* strains containing these vectors were termed HSL 002, HSL 229 and HSL 230 respectively.

### 2.6 Cultivation of strains

*Cupriavidus necator* H16 was obtained from the German Collection of Microorganisms and Cell Cultures, strain number DSM-428. A PHB-negative mutant strain was also used, DSM-541 (Raberg et al., 2014). Cells were cultivated on complete (LB) medium, or minimal medium depending on experimental setup. Minimal medium was composed of 0.78 g/L NaH_2_PO_4_, 4.18 g/L Na_2_HPO_4_x2H_2_O, 1 g/L NH_4_Cl, 0.1 g/L K_2_SO_4_, 0.1 g/L MgCl_2_x6H_2_O, 1.6 mg/L FeCl_3_x6H_2_O, 0.4 mg/L CaCl_2_, 0.05 mg/L CoCl_2_x6H_2_O, 1.8 mg/L Na_2_MoO_4_x2H_2_O, 0.13 g/L Ni_2_SO_4_x6H_2_O, 0.07 mg/L CuCl_2_x2H_2_O. As the carbon source either 2 g/L fructose or 2 g/L pH-neutralized formic acid was added. All cultures were grown at 30 °C. Strains of *E. coli* were grown on LB medium. To overcome the innate resistance of *C. necator* to kanamycin, strains containing pBBR1 based plasmids were cultivated at 150 µg/mL kanamycin.

### 2.7 Assay of AclAB enzyme activity in cell extracts

Ten mL cultures of *E. coli* or *C. necator* AclAB positive clones were grown overnight at 37 °C or 30 °C to approximately OD_600_ 1 and harvested by centrifugation at 4000 g for 5 min. Cell pellets were resuspended in 100 µL of lysis buffer (150 mM NaCl, 50 mM Tris, 1% Triton, pH 8) and lysed using six cycles of bead beating (motor speed 6.5 m/s for 30 s) on an IKA ULTRATURRAX T-25 homogenizer, with 30 s incubation on ice between cycles. Samples were centrifuged at 12000 g for 2 min and the supernatant was collected. The concentration of protein in the supernatant was calculated using a Bradford assay. The AclAB enzyme assay was modified slightly from (Kim and Tabita, 2006). A 1x enzyme mastermix was prepared containing 100 mM Tris-Hcl (pH 8.4), 20 mM sodium citrate, 10 mM dithiothreitol, 10 mM MgCl2, 0.25 mM NADH, 3.3 U of malate dehydrogenase, 0.44 mM CoA and 2.5 mM ATP. Lysate protein (5 µg) was added to 50 µL of mastermix and made up to a final volume of 60 µL with 100 mM Tris-Hcl buffer (pH8.4) in a 96 well plate. The reaction was followed by the decrease in NADH (change in Abs at 340 nm) over 30 min using a TECAN plate reader.

### 2.8 Detection of Xfpk in cell extracts by Western blot

Protein was harvested and quantified using the same methods as for the aclAB enzyme assay above and 5 µg of protein was run on a stain free protein gel (Biorad). The Xfpk protein was visualized using anti-His antibodies HIS.H8 MA1-21315 (Invitrogen) coupled with a WesternBreeze chromogenic immunodetection kit (Invitrogen).

### 2.9 PHB quantification

PHB was quantified in all strains using a modified version of the method developed by (Law & Slepecky, 1961). Samples were pre-cultured in minimal medium with 2 g/L fructose with strain appropriate antibiotics prior to transfer to minimal medium containing formate. Cultivation occurred under either carbon or nitrogen limiting conditions on 2 g/L fructose and 2 g/L formate with strain appropriate antibiotics. To compensate for the lower cell mass obtained from formate, 500 mL cultures were prepared, compared to the 50 mL cultures used for fructose. Concentration of added ammonia required for N limitation was determined experimentally to be 0.1 and 0.01 g/L of ammonium chloride for 2 g/L fructose and formate respectively. For C-limited cultures 1 g/L ammonium chloride was used. Cultures were induced with 1 mM arabinose upon inoculation in the C- or N-limiting media. After 92 h, the 50 mL (fructose) or 500 mL (formate) culture volume was harvested via centrifugation at 4000 g, 20 min, and the supernatant was discarded. Cell pellets were washed in deionized water, transferred to empty, pre-weighed Eppendorf tubes, and incubated at 50 °C for 4 h for liquid evaporation. Tubes were weighed to determine the dry cell mass. Dry cell pellets were dissolved in 1 mL 14% sodium hypochlorite solution and incubated at 37 °C for 1 h with frequent vortexing to aid homogenisation. After incubation, samples were centrifuged at 16000 g for 5 min. The supernatant was removed and the pellets were washed sequentially in 1 mL deionized water, 1 mL acetone, and 1 mL neat ethanol. Between each wash, samples were centrifuged at 16000 xg for 5 min and the supernatant was removed. After the ethanol wash, pellets were resuspended in 1 mL chloroform and transferred to glass tubes. The tubes were incubated at 70 °C for 2 min on a heat block to extract the PHB from the sample. The supernatant was transferred to a second glass tube and the extraction was repeated with 1 mL of chloroform. The 2 mL sample was centrifuged at 4000 g for 5 min to pellet any debris before the chloroform samples were filtered through a 0.22 µm PTFE syringe filter (Whatman). The flow-through was left to evaporate to dryness on a heat block at 37 °C in a fume hood. Upon evaporation the PHB in the samples formed a layer covering the inside of the glass tube. The PHB film was hydrolyzed to crotonic acid by addition of 1 ml of concentrated sulphuric acid and tubes were incubated at 100 °C for 20 min. The resulting crotonic acid samples were diluted 1:100 with 14 mM sulphuric acid and the absorbance at 235 nm was measured (Law & Slepecky, 1961) The amount of PHB in the samples was calculated from a standard curve, prepared from 100 mg sodium 3-hydroxybutyrate hydrolyzed to crotonic acid with concentrated sulphuric acid and diluted 1:100 in 14 mM sulphuric acid. To reduce the effect of any background absorbance values from substances other than crotonic acid, a single sample of strain DSM541 was grown on 2 g/L fructose under C-limited conditions. This sample was subjected to the same extraction protocol as described above and its absorbance value at 235 nm was deducted from all other samples.

## 3. Results and discussion

### 3.1 Network flexibility in *C. necator* is high on fructose and low on formate

We began by enumerating all EFMs in the core network connecting fructose or formate to biomass or PHB and subjected the EFMs to MDF analysis, which determined the highest driving force of the least thermodynamically favorable reaction of the EFM (Figure 1A). If the calculated MDF was negative, that EFM was considered thermodynamically infeasible and excluded from further analysis (Figure 1B). Reactions were to a large extent set to be reversible, with exceptions such as exchange reactions and reactions known to operate only in the forward direction, such as RUBISCO. Avoiding reaction direction constraints prior to thermodynamic analysis was recently shown to be crucial in finding carbon fixation pathways in *E. coli* (Satanowski et al., 2020). The total number of feasible EFMs leading to either biomass, PHB, or both, can be taken as a measure for the flexibility of the metabolic network. High flexibility was found on fructose, resulting in 9705 feasible EFMs (Figure S1). We note that CBB cycle flux was allowed when utilizing fructose, which increased flexibility by allowing potential recapture of CO_2_ released by decarboxylation reactions. On formate, only 396 thermodynamically feasible EFMs were found. Formate is assimilated only *via* the CBB cycle, so metabolic flexibility was reduced and the network was more rigid. The rigidity of the network can be estimated from the fraction of EFMs that use a certain reaction. If a reaction was used in a majority of the EFMs, the network is more rigid in that dimension. Reactions used in all EFMs are considered essential. On formate, RUBISCO and PRK were essential and other CBB cycle reactions such as RPE, RPI, TKT, PGI, and PGK had forced directionality (Figure S2). The reversal of the reaction of the soluble transhydrogenase STH was also forced, so that NADPH (required for CBB cycle reactions and PHB synthesis) was produced from NADH derived from formate. Fructose is metabolized *via* the glycolytic ED pathway (Pohlmann et al., 2006), and these had high usage (Figure S2).

Two heterologous enzymes were considered for their ability to affect PHB production (Figure 1C). The promiscuous enzyme phosphoketolase (Xfpk) would form a shortcut from the PPP/CBB cycle towards acetyl-CoA *via* formation of acetyl-phosphate from F6P or X5P (XFPK reaction; (Bogorad et al., 2013)). Using XFPK reactions instead of the canonical lower glycolytic route to acetyl-CoA circumvents the decarboxylation reaction of pyruvate dehydrogenase (PDH). Expression of Xfpk enzyme increased production of acetyl-CoA derived products in heterotrophic and autotrophic microorganisms (Anfelt et al., 2015; Kocharin et al., 2013; Qin et al., 2020). Studies on tri-functional Xfpk, acting on X5P, F6P, and S7P, showed that its activity can alter flux distributions in the PPP, allowing TKT to be knocked out (Hellgren et al., 2020; Krüsemann et al., 2018). *C. necator* does not possess an Xfpk, though a heterologous Xfpk was expressed previously and could rescue growth on fructose upon inactivation of the ED-pathway (Fleige et al., 2011). The enzyme ATP-citrate lyase (AclAB), also absent in *C. necator,* catalyzes the ATP-dependent cleavage of citrate into acetyl-CoA and oxaloacetate (ACL reaction). The resulting decrease in net flux towards the TCA cycle is envisioned to provide an increased acetyl-CoA pool available for PHB production. A reduced flux through citrate synthase increased the specific productivity of acetyl-CoA derived products such as propanol and n-butanol in engineered bacteria (Shabestary et al., 2018, 2021; Soma et al., 2014). Addition of the two primary XFPK reactions (F6P and X5P cleavage, respectively) increased network flexibility drastically, adding 168139 EFMs and 4392 EFMs for fructose and formate, respectively (Figure S1). Addition of the XFPK reactions also reduced network rigidity, as evidenced by reduced usage of PDH in these EFMs (Figure S2). Similarly, usage of RPE and TKT1 was also reduced, showing how XFPK reactions can form an effective shortcut from the PPP/CBB cycle. EFMs using ACL had fewer options to supply the relevant reactant (citrate) and therefore few additional EFMs were found (Figure S1). In fact, no feasible EFMs to biomass or PHB using the AclAB reaction were found on formate. However, we note that flux distributions could exist that have non-zero flux through ACL, through combination with EFMs that have byproduct formation. The addition of ACL had negligible effect on network rigidity (Figure S2).

### 3.2 Metabolic strategies defined by thermodynamic limits and yields

For each EFM we determined the biomass yield, the PHB yield, and the MDF value (Figure 2A-B, Figure S3). While the maximum biomass yield observed for formate (∼0.12 g_biomass_/g_substrate_) corresponded well to experimental results (Grunwald et al., 2015; Jahn et al., 2021), the model overestimated the maximum biomass yield for fructose (∼0.53 g_biomass_/g_substrate_, Figure 2A). *In vivo* flux distributions on this substrate are therefore likely to contain EFMs with lower yields. The overall lower biomass and PHB yields on formate are due to its lower degree of reduction compared to fructose, as well as the energetic demands of the CBB cycle. The energy of three formate molecules is required to fix one molecule of CO_2_ via the CBB cycle (Grunwald et al., 2015). Synthetic formate assimilation pathways provide the potential to increase biomass yield on formate (Claassens et al., 2020), (Löwe & Kremling, 2021), though these were not considered here. All enumerated EFMs fell within the biomass-PHB production envelope (Figure 2A-B, shaded area), which contains every possible flux distribution to these products that can be obtained by flux balance analysis (FBA). Many EFMs lead exclusively to either biomass or PHB production (points occupying the x- or y-axis in Figure 2A-B). For formate, no EFM supported biomass and PHB production simultaneously, suggesting an orthogonality between growth and PHB production on this substrate.

The MDF value of an EFM represents the lowest driving force and therefore is an approximation of the highest metabolic burden to enable a certain flux, as more enzymatic activity is “wasted” on the reverse reaction if driving force is low (Noor et al., 2014). EFMs with the same yield or MDF suggest that a common “metabolic strategy” is employed among them, such as a common reaction or set of reactions that constrains stoichiometry or overall thermodynamics. Our detection of the same MDF value in multiple EFMs for both substrates suggests the presence of global thermodynamic bottlenecks (Figure 2C-D). Nonetheless, the higher metabolic flexibility on fructose allowed the existence of metabolic strategies with MDF values of up to ∼13.2 kJ/mol, compared to ∼5.4 kJ/mol for formate. This implied that utilizing fructose could circumvent thermodynamic limitations found for formate utilization.

The addition of XFPK reactions increased the maximum attainable yields for PHB on both fructose and formate, and biomass yield on formate, by circumventing the decarboxylation of pyruvate *via* PDH. Addition of XFPK reactions also increased the maximum MDF for biomass on fructose, from ∼10 kJ/mol to ∼13.6 kJ/mol, but did not change the MDF on formate (Figure S3). EFMs using ACL did not increase the maximum attainable MDF or maximum yield; the production envelope of the ACL-containing network had the same shape as the wild type. Further analysis revealed that forcing flux through ACL was always associated with a reduction in yield compared to the FBA optimum. With the citrate pool always being supplied from acetyl-CoA, ACL usage forms a cyclic pathway reducing maximum yields but not necessarily violating thermodynamic constraints due to the associated ATP-hydrolysis. Although no yield improvements via ACL were observed, increased driving forces towards acetyl-CoA were nonetheless possible (section 3.5).

### 3.3 Identifying reactions that define the yields and thermodynamics of metabolic strategies

We next sought to analyze what role *individual* reactions play in determining the yield and MDF of EFMs. We devised a quantitative scoring system for whether a particular reaction is associated with EFMs that have higher yields or MDF values than average (Methods, 2.3). In this scoring system, a reaction having a biomass yield score of 10 means that the average biomass yield of EFMs using this reaction is 10% higher than the average biomass yield over all EFMs (Supplementary Tables S1-2 for all reaction scores and p-values). An example interpretation of the scoring plots shown in Figure 3 is the negative impact of pyruvate excretion on both biomass and PHB yield when fructose is utilized. Carbon channeled into pyruvate to be excreted will not participate in biomass or PHB, resulting in reduced yield for both.

**Figure 3:**
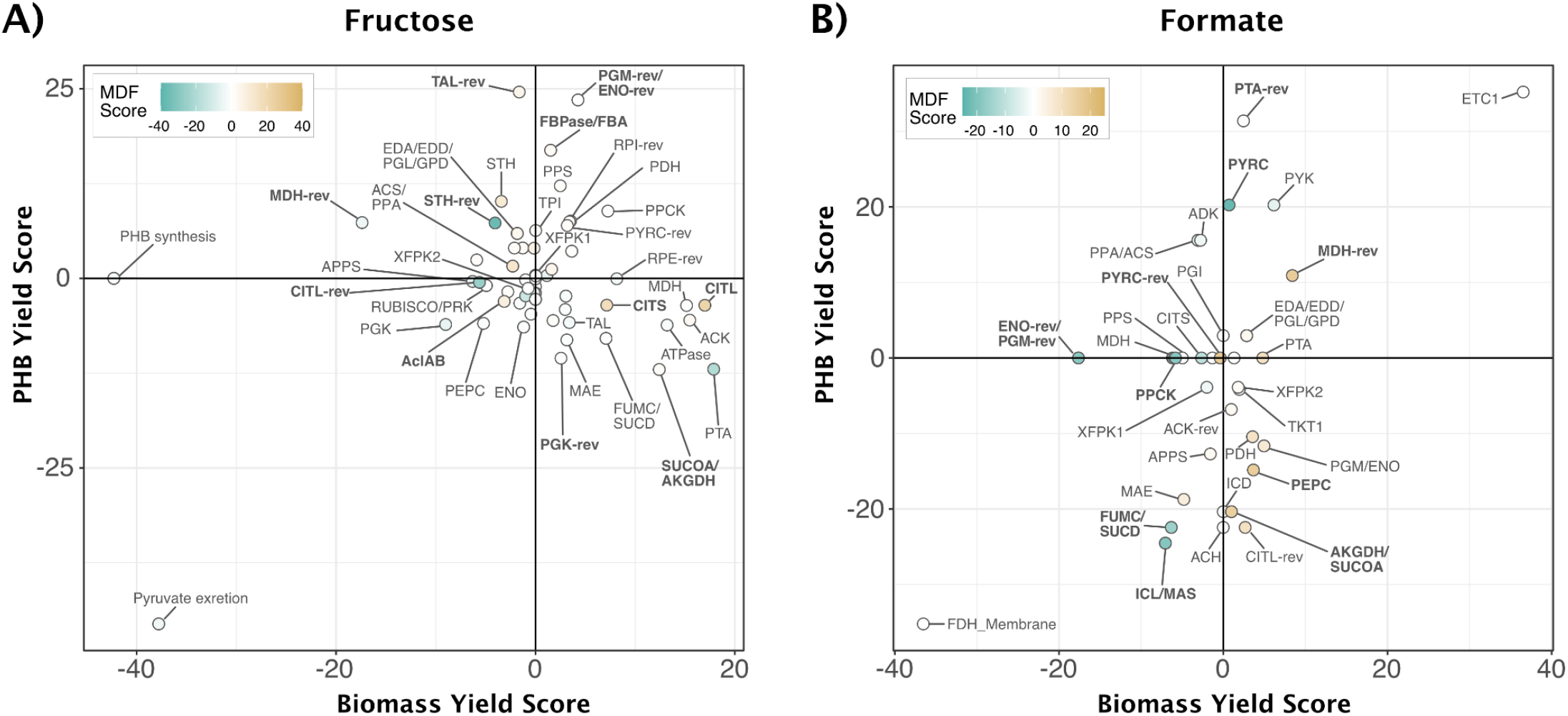
Influence of individual reactions on biomass yield and MDF. **A)** Biomass-, PHB- and MDF scores for individual reactions, for fructose. Scores indicate positive and negative influence on yields or MDF. Reactions participating in all EFMs would accumulate at the origin and were therefore excluded from the figure. **B)** Same as in A) but for formate. Only the reactions participating in both biomass and PHB production are shown (see Supplementary Tables S1-S2 for all scores).

When considering potential targets for metabolic engineering to improve PHB yields, reactions with negative PHB yield scores are of highest interest (bottom quadrants in Figure 3), since blocking or reducing their activity could be beneficial. Reactions in the bottom right quadrants were negatively associated with PHB production, and also had a positive effect on biomass production. Some targets in this quadrant were common for both fructose and formate, including TCA cycle reactions AKGDH, SUCOA, FUMC and SUCD. Targeting these reactions for repression could channel carbon away from biomass and towards PHB, decoupling product formation and growth (Pandit et al., 2017). A recent study on production of 1,3-butanediol (derived from acetyl-CoA) by *C. necator* showed that knockout of the SUCD, increased titers of 1,3-butanediol 2-fold when grown on gluconate. Though SUCD had a positive biomass yield score, there was no apparent effect on growth rate or biomass yield (“Engineering Cupriavidus Necator H16 for the Autotrophic Production of (R)-1,3-Butanediol,” 2021).

There were some substrate-specific trends. On formate, a functional TCA cycle and glyoxylate shunt scored lowest regarding PHB yield, while high yield scores were found for reactions around acetyl-CoA metabolism, including PYRC (pyruvate carboxylase), ACS (acetyl-CoA synthetase), ADK (adenylate kinase), PPA (pyrophosphatase) and the reversal of PTA (to form acetyl-CoA from acetyl-phosphate) (Figure 3B). Due to the low number of EFMs on formate, p-values were high and the results should be interpreted with care (Supplementary Table S2). Conversion of acetyl-P to acetyl-CoA (or acetyl-P to acetate to acetyl-CoA) is part of the high-yield Xfpk route from sugar phosphates to acetyl-P (Figure 1). However, Xfpk did not have a high yield score. This apparent disagreement arises because the addition of Xfpk allows both high and lower-yield EFMs, so its yield score is not high. On fructose, a functional CBB cycle, represented by PRK and RUBISCO, had a negative score for biomass yield, which agrees with yields determined by proteomics-constrained flux simulations (Jahn et al., 2021). The CBB cycle requires ATP and NADPH, whose production requires decarboxylation in lower glycolysis and the TCA cycle, diverting carbon away from biomass. An active CBB cycle in *C. necator* does allow for a higher yield of PHB on fructose (Shimizu et al., 2015), though yield scores for PRK and RUBISCO were neutral relative to average PHB yields.

Many reactions had negligible MDF scores, emphasizing that the thermodynamic limitations of EFMs are defined by only a few impactful reactions. Highly positive MDF scores were found for CITL and CITS on fructose, and for PEPC, AKGDH, SUCOA and reverse reactions of MDH and PYRC on formate. Together with MAE, reversals of PYRC and MDH form a cycle, where PYR is carboxylated to OAA with hydrolysis of ATP, reduced to malate and decarboxylated to pyruvate. This thermodynamically efficient shunt would avoid PYRC and MDH, which had low MDF scores, at a cost of one ATP and no net production of NADH.

Which EFMs are preferentially used *in vivo* could be better predicted with additional physiological constraints on the solution space, such as protein abundance, gene essentiality, and enzyme kinetics and biochemical regulations (Gerosa et al., 2015; Jahn et al., 2021; Sánchez et al., 2017). Along those lines, a transposon-library of *C. necator* identified ACS and PEPC as essential reactions, hinting towards their usage *in vivo* (Jahn et al., 2021).

### 3.4 Metabolic strategies toward PHB have driving forces limited by a few common reactions

For each EFM, the MDF algorithm finds a set of metabolite concentrations that maximizes the thermodynamically least favorable reaction; the calculated driving force of this reaction is the MDF (maximized minimum driving force). Knowledge of the MDF, as well as *which* reaction in the EFM operates at the MDF, is useful for metabolic engineering, as reactions with low driving forces may require more enzyme resources to achieve a high net flux. However, the MDF solution is not unique; there can be many combinations of metabolite concentrations resulting in the same MDF, as well as multiple reactions operating at the MDF. This variability makes it difficult to identify precisely which reactions are likely thermodynamic bottlenecks. In this section, we sought to mitigate this degeneracy problem by first restricting possible metabolite concentration ranges, followed by randomly sampling metabolite concentrations within those ranges and calculating reaction driving forces. The sampling of metabolite concentrations (and thus driving forces) allows an estimate of the probability for a given reaction to have a certain driving force.

To determine metabolite concentration ranges, we performed a metabolite variability analysis for each EFM. First, the MDF of the EFM was determined. Next, metabolite concentrations were varied to determine which ranges of metabolite concentrations still allowed the MDF (down to 90% of this optimal value) to be achieved. Metabolites with a narrow allowable metabolite concentration range were considered to be likely involved in the MDF reaction. The global, substrate-independent metabolic strategies were constrained by the same metabolites. For example, for all EFMs with an MDF of ∼4.6 kJ/mol, the concentrations of citrate, acetate, isocitrate and oxaloacetate were tightly constrained, on both fructose and formate (Figures S4-S5). In other metabolic strategies, these metabolites had broader concentration ranges, or were constrained to a different value (see for example oxaloacetate (OAA) in Figure S4). Generally, a higher MDF resulted in narrower concentration ranges for many metabolites (Figures S4).

Narrow concentration ranges around an MDF could be indicative of which reactions were thermodynamically limiting, but not conclusive. To get a probabilistic estimate of a reaction’s likelihood of being the MDF, we estimated the driving force of each reaction in all EFMs by sampling metabolite concentrations for that EFM, constrained by the MDF as described above. Figure 4A shows selected reactions that had a driving force equal to the MDF value or close to it, and thereby marked the thermodynamic limit (see Figure S6 for details, and figures S7-S8 for all reactions). Often, several reactions operated at the MDF value, implying that combinations of reactions or subnetworks determined the EFM thermodynamics. Metabolic strategies common to both substrates shared the same limiting reactions, most prominently the citrate-acting reactions ACH and reverse of CITL for MDF values at ∼4.6 kJ/mol (Figure 4A). The reverse CITL reaction was always limiting when it was used, which suggests a high enzyme expression level if this metabolic strategy were to be employed *in vivo* (Noor et al., 2014). Interestingly, a high expression of citrate lyase subunit *CitE* was reported in PHB-negative strains of *C. necator*, were it was suspected to be involved in the forward direction for anaplerotic recycling of citrate to oxaloacetate and acetate (Raberg et al., 2014). ACH and MDH, identified as thermodynamically limiting reactions by our analysis, also showed high enzyme levels in a recent study on proteomic adaptations in *C. necator* (Jahn et al., 2021). While operation closer to equilibrium may require more enzyme resources to ensure a net flux, it also affords a faster switch in reaction directionality and moves control of the associated reaction rates to the thermodynamic level, i.e. the relationship between the substrate and product concentrations (Park et al., 2019), rather than by expression of the catalyzing enzyme. A higher DF and flux control at the level of enzyme expression is more amenable to engineering, but could render metabolism less robust against sudden environmental fluctuations such as changes in substrate (Basan et al., 2020). While these dynamics are beyond the scope of this study, identifying metabolic strategies at steady-state can help elucidate the switches necessary to adapt to certain conditions (Gerosa et al., 2015).

**Figure 4:**
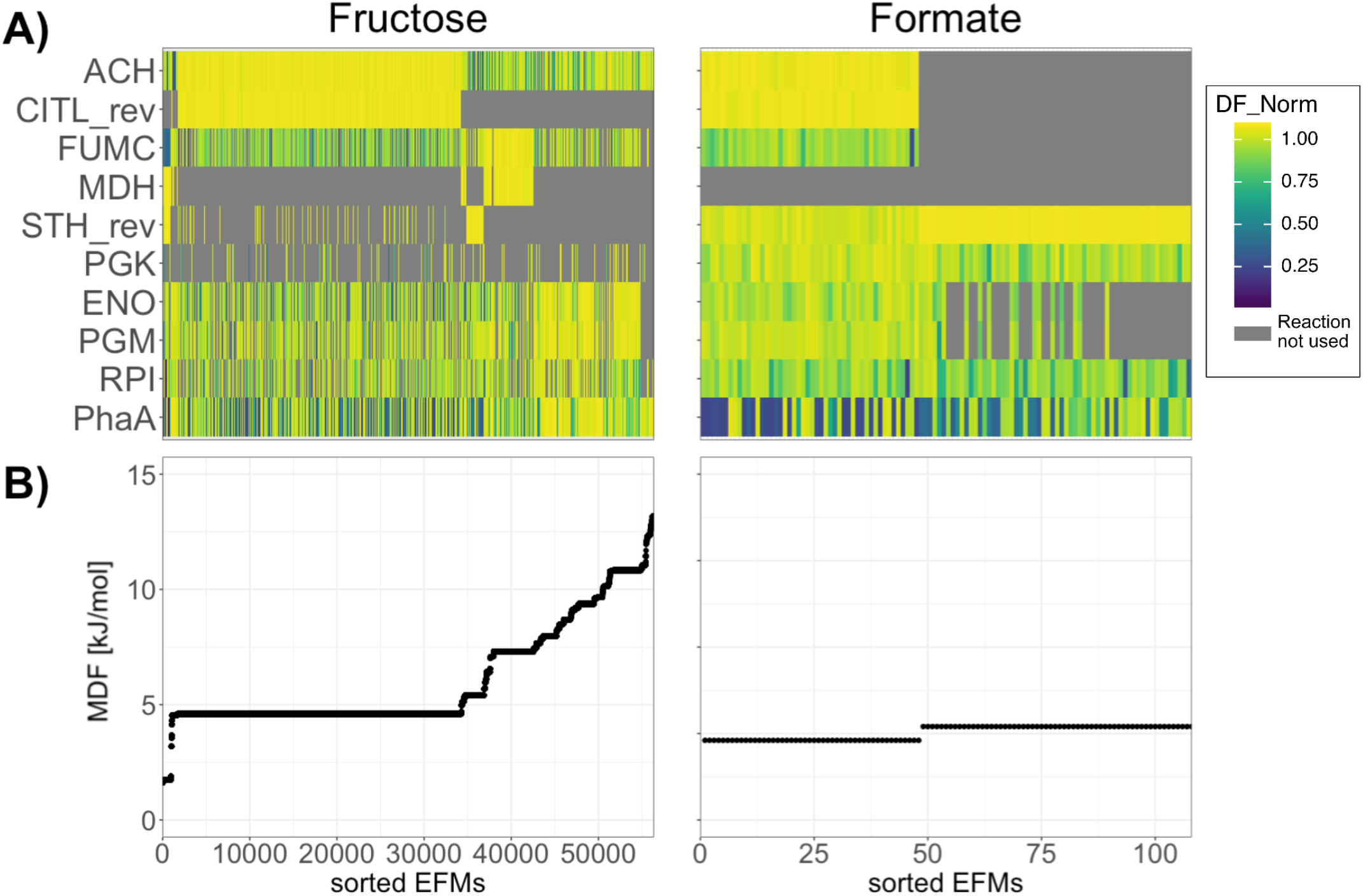
Thermodynamically limiting reactions for PHB production identified from metabolite sampling. For each EFM, metabolite concentrations were sampled and DF calculated. **A)** Heatmaps showing relative closeness of each reaction to the MDF. The higher the normalized DF, the closer that reaction’s DF is to the MDF. Each column represents an EFM. EFMs are sorted first by their MDF and in case of degenerate MDF, by the mean of all DFs. Yellow: this reaction operates at the MDF. Gray: This reaction is not used in this EFM. **B)** MDFs of EFMs, sorted by MDF value.

Finally, we sought to explore the thermodynamic landscape for PHB production from formate as one of the most promising renewable feedstocks for bioprocesses (Claassens et al., 2019). Two metabolic strategies, *i.e.* multiple EFMs with the same MDF, were identified for PHB production on formate (Figure 2D, Figure 4B). Plotting the mean DF over all reactions used in EFMs of the two separate strategies revealed reaction usage and limiting steps (Figure 5). In metabolic strategy 1 (MDF = 4.6 kJ/mol) the TCA cycle was used, and the first two steps (CITL_rev and ACH) marked the thermodynamic limitation for all EFMs of this metabolic strategy. In metabolic strategy 2 (MDF = 5.4 kJ/mol), the ED pathway was used, but the reaction constraining the MDF for all EFMs of that strategy was identified as the transhydrogenase (STH_rev), marking the cofactor balance as the thermodynamic limitation. In both strategies, several reactions had mean DFs close to the MDF values, including lower glycolysis, as well as the ATP-consuming reaction PGK and the “carbon shuffle,” in the CBB cycle. As mentioned above, operating closer to thermodynamic equilibrium moves the flux control to the metabolite levels (Park et al., 2019), while high DFs indicate control at the expression level. Common reactions with high DFs in the two metabolic strategies were found in the anaplerotic reactions (MAE, PEPC), as well as some CBB cycle reactions (RUBISCO, PRK, GADPH, FBA, FBPase, SBPase), implying lower expression levels to carry flux. However, having a high DF is no guarantee of low expression levels, as is exemplified by the high protein levels of RUBISCO, an enzyme limited by its low rate (Jahn et al., 2021). While it is challenging to precisely determine which metabolic strategy (or combination of EFMs) is used for PHB production *in vivo* based on stoichiometry and thermodynamics alone, essentiality analysis, such as through use of genetic knockout libraries (Jahn et al., 2021), could help to elucidate the actual *in vivo* flux distribution to PHB.

**Figure 5:**
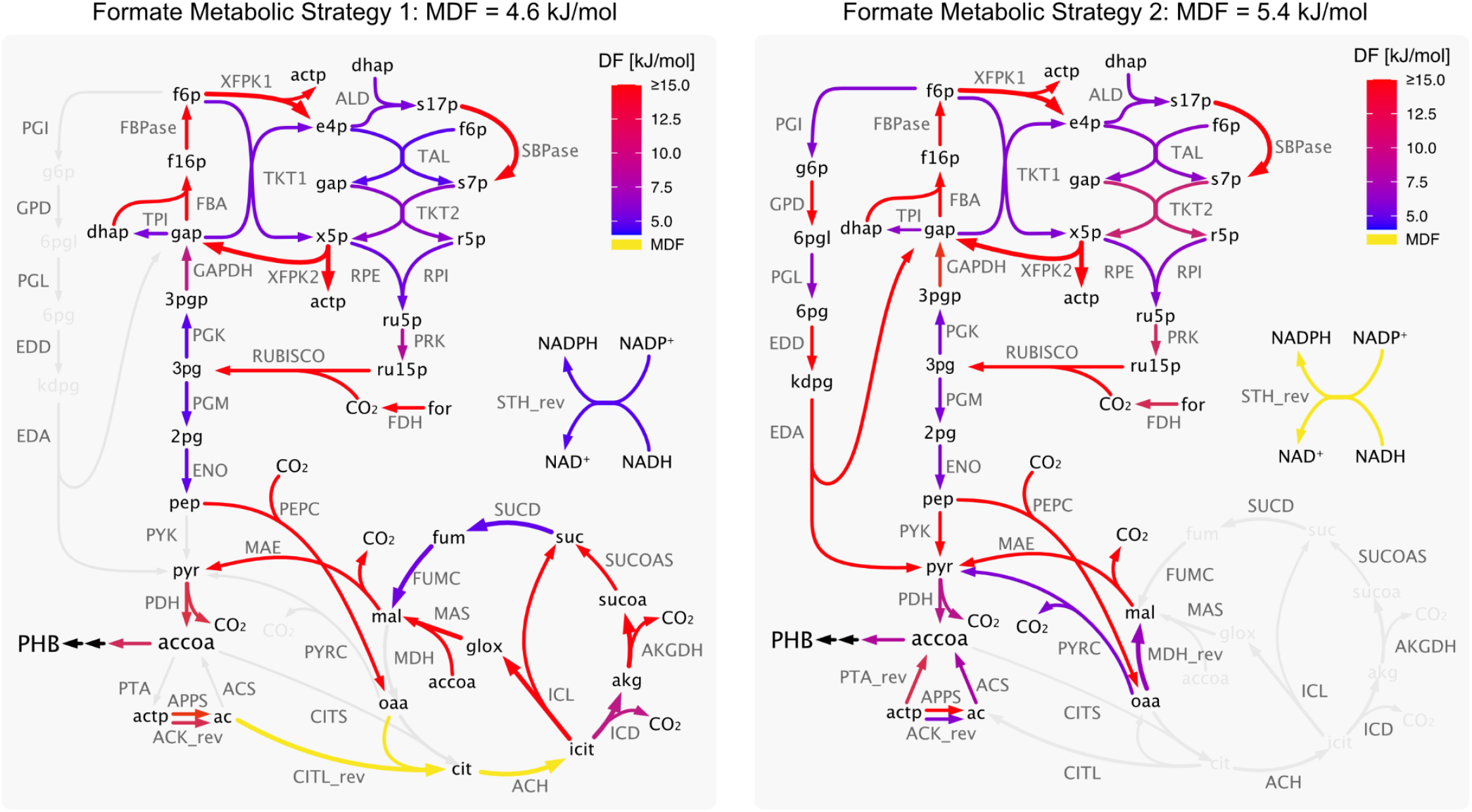
Reaction driving forces for two metabolic strategies to PHB from formate. Metabolic network maps showing mean DF from random sampling over all EFMs for the two metabolic strategies found for formate. Reactions marked in yellow indicate reactions operating at the MDF value for all EFMs of that metabolic strategy.

### 3.5 Xfpk and AclAB increase PHB production by affecting driving force or yield

We showed previously that addition of XFPK reactions to the network increased maximum attainable yields, and allowed EFMs with higher driving forces, while addition of ACL did not (Figure 2). However, it is possible that ACL could impart a higher DF to reactions closer to PHB, and thus influence metabolic rates to this product. We used the results from the metabolite sampling algorithm to calculate the median driving forces of reactions producing acetyl-CoA for each EFM. The comparison revealed that both the addition of ACL and XFPK reactions significantly increased the driving forces towards acetyl-CoA on fructose (Figure 6A). Addition of XFPK also significantly increased driving force towards acetyl-CoA on formate (Figure 6B). Since ACL was not used by any EFM on formate, its effect was not evaluated on this substrate.

**Figure 6.**
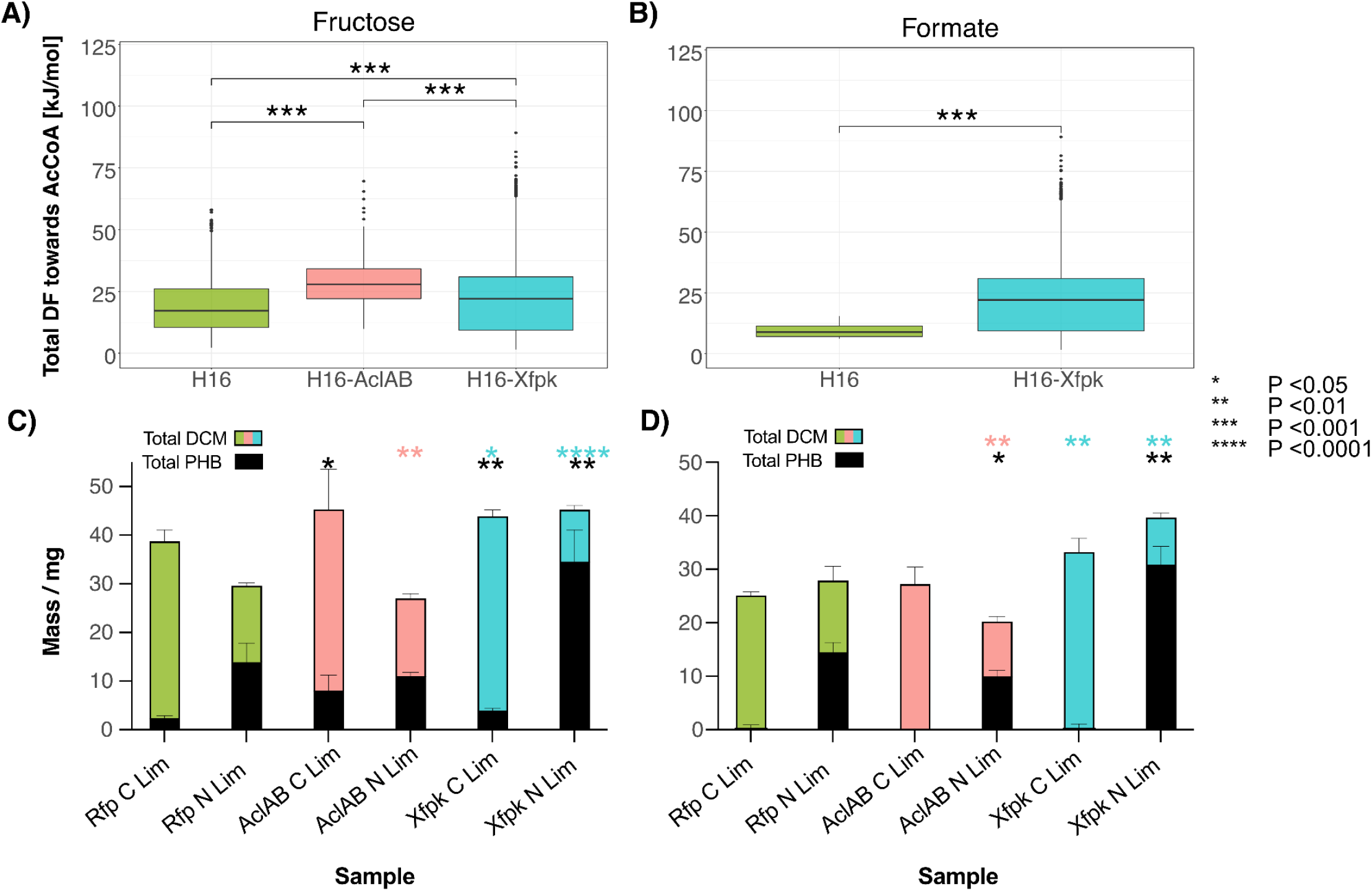
Driving forces toward acetyl-CoA and PHB content of mutant strains. **A)** Boxplots for sums of calculated driving forces of all reactions leading towards acetyl-CoA for each EFM for fructose, for each substrate and strain. Significance (Wilcoxon rank sum test) displayed. **B)** Same as A) but for formate. No EFMs using AclAB were found. **C)** Average total dry cell mass (DCM) and mass of PHB of cultures grown in C or N limited conditions on fructose. Significant differences in DCM and PHB are indicated by * in the color corresponding to the strain and black, respectively. **D)** Total dry cell mass (DCM) and mass of PHB of cultures grown in C or N limited conditions on formate. Significant differences in DCM and PHB are indicated by * in the color of the corresponding strain and black, respectively. Statistics calculated using students T-test and comparison to Rfp cultures, n = 3.

To test model predictions on the effect of Xfpk and AclAB on PHB production, we created *C. necator* strains expressing either Xfpk enzyme from *Bifidobacterium breve* or AclAB enzyme from *Chlorobium limicola*. A strain expressing Rfp was created to use as a control. The expression of the heterologous enzymes was confirmed by measured enzyme activity in cell lysates (AclAB) (Figure S9), visually (Rfp) or western blot (Xfpk) (Figure S10). These strains were screened for PHB accumulation on both fructose and formate in both carbon-limited and nitrogen-limited batch cultures (Methods). In the C-limited cultures, cells grew until the carbon source was exhausted. In N-limited cultures, cells were grown to stationary phase and left for an additional 48 hours to allow the residual carbon to be utilized for PHB production.

The Xfpk strain accumulated 60% more PHB than the control strain during N-limitation on both fructose and formate (Figure 6C-D, Table 1), with PHB increasing from an average of 478 and 422 mg PHB/gDCM to an average of 766 and 778 mg PHB/gDCM, respectively (Figure 6C and 6D). During C-limitation on fructose, PHB accumulation was detectable at low levels in both strains with a small but statistically significant increase from an average of 65 to 95 mg PHB/gDCM. On formate, no PHB accumulation was detected during C-limitation in any strain. This is in line with model results that no EFMs on formate produce both biomass and PHB, while many EFMs on fructose lead to both products (Figure 2). The Xfpk strain showed an increase in total DCM yield on both fructose and formate, which ranged from 15% (fructose) to 30% (formate). This increase in yield is a general confirmation of model predictions that usage of Xfpk allows higher biomass yields (Figure 2). The AclAB strain did not accumulate more PHB than the control strain, except for during C-limited growth on fructose. In this condition, the AclAB strain accumulated an average of 191 mg PHB/gDCW, more than three times that of the control strain. The increase in PHB content may be a reflection of a “forced” accumulation of acetyl-CoA by the ATP-driven cleavage of citrate. AclAB did not affect biomass yields on fructose, and reduced biomass yields on formate.

**Table 1.** Total dry cell mass (DCM), total mass of PHB and PHB as a percentage of total mass for strains under carbon (C) and nitrogen (N) limitation on fructose and formate. Values are averages of 3 measurements +/- standard deviation

|  |  | **Rfp** |  | **AcIAB** |  | **Xfpk** |  |
| --- | --- | --- | --- | --- | --- | --- | --- |
|  |  | **C-limited** | **N-limited** | **C-limited** | **N-limited** | **C-limited** | **N-limited** |
| **Fructose** | **DCM (mg)** | 39.0 +/- 2.1 | 29.9 +/- 0.3 | 45.5 +/- 8.0 | 27.2 +/- 0.8 | 44.1 +/- 1.1 | 45.5 +/- 0.7 |
|  | **PHB (mg)** | 2.6 +/- 0.3 | 14.3 +/- 3.5 | 8.3 +/- 3.0 | 11.2 +/- 0.6 | 4.2 +/- 0.2 | 34.8 +/- 6.2 |
|  | **Ratio (%PHB)** | 6.6 | 47.8 | 18.2 | 41.2 | 9.5 | 76.5 |
| **Formate** | **DCM (mg)** | 25.3 +/- 0.5 | 28.1 +/- 2.5 | 27.4 +/- 3.0 | 20.4 +/- 0.8 | 33.4 +/- 2.3 | 39.9 +/- 0.6 |
|  | **PHB (mg)** | 0.3 +/- 0.6 | 14.7 +/- 1.7 | 0 +/- 0 | 10.2 +/- 1 | 0.4 +/- 0.6 | 31.1 +/- 3.2 |
|  | **Ratio (%PHB)** | 1.2 | 52.3 | 0 | 50 | 1.2 | 77.9 |

## 4. Conclusion

This study shed light on the yields and driving forces of natural and engineered metabolic strategies for PHB production in *C. necator*. We considered fructose and formate, which are metabolized by different pathways and have different carbon and energy contents, as well as augmenting wild-type metabolism with two heterologous reactions. In this context, we enumerated EFMs (Terzer & Stelling, 2008) and subsequently used the MDF algorithm (Noor et al., 2014) to judge thermodynamic optimality, and compared each reaction’s influence on yields and MDF. The yield and MDF scores are a way to extract useful information from thousands of EFMs, for example highlighting genes that are associated with EFMs of low yield or driving force. These genes could be suppressed in two-stage cultivations with dedicated growth and production phases.

The use of EFMs allowed us to explore the entire solution space, which is crucial as the *in vivo* flux distributions are often not known and may not follow the optimality criteria used for methods such as FBA. One significant insight from this work is that many EFMs have the same MDF value, and these MDFs are determined by a few reactions and metabolites on both fructose and formate. Most EFMs had an MDF value of about ∼4.6 kJ/mol, and were limited by ACH and CITL reactions, which are connected *via* tightly constrained concentration ranges of citrate, isocitrate, oxaloacetate and acetate. Changes in the concentrations of these metabolites could therefore be controlling factors *in vivo*. The thermodynamic driving force gives insight into how flux through a certain reaction could be regulated; low driving force makes reaction directionality and flux susceptible to changes in metabolite concentrations. Such reactions may therefore be more flexible in directionality depending on substrate. Engineering strategies should therefore be aimed at adjacent reactions which influence the metabolite concentrations around the reaction with low driving force.

This work also provides an additional tool for MDF analysis, particularly of EFMs. Sampling of metabolite concentrations (and driving forces) is a way to visualize the probable driving forces of all reactions in an EFM. While we restricted our analysis to fructose and formate, other substrates can be easily integrated, such as glycerol. Glycerol could be considered a mixotrophic substrate, where the CBB cycle may be active (Alagesan, Minton, et al., 2018). Furthermore, our methodology could be applied to metabolic networks with additional heterologous reactions, such as novel formate assimilation pathways recently implemented in *C. necator* (Alagesan, Minton, et al., 2018; Claassens et al., 2020).

The different effects on PHB production of Xfpk and AclAB on fructose and formate are illuminating for future metabolic engineering efforts. When developing engineering strategies it is important to consider the outcome that is to be optimized. During C-limited growth on fructose, expression of AclAB led to an increase in PHB, presumably by increasing driving force to acetyl-CoA, even though yield was not increased. It is crucial to consider also the effect of substrate, as this increased DF resulted in a decrease in both biomass and PHB production during growth on formate. The forced flux due to expression of AclAB resulted in lower yield flux distributions. This reduction in growth is in line with the severe restriction in growth strategies seen on formate compared to fructose, whereby no viable AclAB utilizing EFMs were seen for PHB production, rendering its expression an energy burden under these conditions (Figure 2). Nevertheless, these results do highlight the benefit of MDF analysis for the development of engineering strategies that prioritize increased DF towards the product at the expense of overall yield.

Metabolic modeling studies always depend on the input data and assumptions used to capture physiological behavior. The MDF algorithm maximizes the driving force of the least favorable reaction, which would minimize that enzyme burden on the cell (Noor et al., 2014). The assumption that a cell would operate at lowest possible enzyme burden is reasonable, but more precise predictions of enzyme abundances would require knowledge of kinetic enzyme parameters and allosteric regulations, as well consideration of incomplete saturation necessary for metabolic robustness against perturbations (Janasch et al., 2019; Sander et al., 2019). Experimental values for uptake and excretion of compounds can additionally aid to constrain the broad solution space resulting from the EFM analysis (Jol et al., 2012). Furthermore, driving forces rely on the Gibbs Free energies of formation of reactants and products, whose estimation is also associated with uncertainties (Noor et al., 2013). Lastly, collecting multi-omics datasets and integrating them with metabolic network analysis will be a necessary next step in understanding the impact of different substrate utilization and genetic perturbations (Donati et al., 2021; Gerosa et al., 2015; Hackett et al., 2016).

## Supplementary Information

### Code availability

All models and scripts for the computational analysis are available at https://github.com/MJanasch/ThermoCup.

### Supplementary Dataset

**Dataset S1** - Reactions and metabolites of the core model of *C. necator*

### Supplementary Methods

**1.** Gibbs energy change of transport reactions between extracellular and intracellular compartments

**2.** MDF framework used to evaluate max-min Driving Force for each EFM

**3.** Strain construction

### Supplementary Figures

**Figure S1 -** Flexibility of *C. necator*’s core metabolism

**Figure S2 -** Reaction usage of *C. necator’s* core metabolism in dependency of the strain and substrate

**Figure S3 -** Relationship of Yields and MDFs for biomass and PHB producing EFMs

**Figure S4 -** Metabolite concentration ranges for EFMs producing PHB under fructose utilization

**Figure S5 -** Metabolite concentration ranges for EFMs producing PHB under formate utilization

**Figure S6 -** Example for derivation of the DF_Norm value

**Figure S7 -** DF_Norm for all reactions for all EFMs producing PHB under fructose utilization

**Figure S8 -** DF_Norm for all reactions for all EFMs producing PHB under formate utilization

**Figure S9 -** AclAB activity assay from *E.coli* and *C. necator* cultures

**Figure S10 -** Western blot of Xfpk from *E.coli* and *C. necator* cultures

**Figure S11 -** Plasmid maps

### Supplementary Tables

**Table S1 -** Biomass and PHB yield scores and MDF scores for all reactions for the case of fructose utilization

**Table S2** - Biomass and PHB yield scores and MDF scores for all reactions for the case of formate utilization

**Table S3 -** Plasmid list

**Table S4** - Strain list

**Table S5** - Primer list

## Acknowledgements

We are grateful to Nuha Salem for discussions in the initial phase of the project. We furthermore thank Nico Claassens and Elad Noor for quick replies to inquiries regarding physiological pH and ionic strength, and EFM enumeration, respectively.

