## Supplementary_Material for "Thermodynamic limitations of metabolic strategies for PHB production from formate and fructose in *Cupriavidus necator*": ThermoCup_Supplements.pdf

for

#### Code availability

All models and scripts for the computational analysis are available at https://github.com/MJanasch/ThermoCup.

#### Supplementary Dataset

**Dataset S1** - Reactions and metabolites of the core model of *C. necator*

#### Supplementary Methods

1. Gibbs energy change of transport reactions between extracellular and intracellular compartments

   Including Table SM1: Concentration ranges for external metabolites that differ from the default range

2. MDF framework used to evaluate max-min Driving Force for each EFM
3. Strain construction

#### Supplementary Figures

**Figure S1** - Flexibility of *C. necator*'s core metabolism

**Figure S2** - Reaction usage of *C. necator's* core metabolism in dependency of the strain and substrate

**Figure S3** - Relationship of Yields and MDFs for biomass and PHB producing EFMs

**Figure S4** - Metabolite concentration ranges for EFMs producing PHB under fructose utilization

**Figure S5** - Metabolite concentration ranges for EFMs producing PHB under formate utilization

**Figure S6** - Example for derivation of the DF_Norm value

**Figure S7** - DF_Norm for all reactions for all EFMs producing PHB under fructose utilization

**Figure S8** - DF_Norm for all reactions for all EFMs producing PHB under formate utilization

**Figure S9** - aclAB activity assay from *E.coli* and *C. necator* cultures

**Figure S10** - Western blot of XFPK from *E.coli* and *C. necator* cultures

**Figure S11** - Plasmid maps

#### Supplementary Tables

**Table S1** - Biomass and PHB yield scores and MDF scores for all reactions for the case of fructose utilization

**Table S2** - Biomass and PHB yield scores and MDF scores for all reactions for the case of formate utilization

**Table S3** - Plasmid list

**Table S4** - Strain list

**Table S5** - Primer list

### Supplementary Methods

#### 1. Gibbs energy change of transport reactions between extracellular and intracellular compartments

The Gibbs energy change associated with reactions of metabolite uptake and excretion was calculated according to Jol et al., 2010 and is detailed in

*ThermoCup_CalculateTransportEnergy.m*

found in the github repository https://github.com/MJanasch/ThermoCup.

Internal pH was set as 7.5, external pH was set as 7. The membrane potential was set to $\Delta\varphi = -0.15$ V, the Faraday constant taken as F ≈ 96.485 C/mmol. The concentration ranges of external metabolites was the same was for internal metabolites ($[m]_{min}$= 1 μM and $[m]_{max}$= 10 mM), with the exceptions shown in Supplementary Methods Table SM1.

Table SM1: Concentration ranges for external metabolites that differ from the default range.

| Metabolite | Name | [m]_min [M] | [m]_max [M] |
|---|---|---|---|
| o2[e] | Oxygen | 5.00E-04 | 5.00E-04 |
| co2tot[e] | total CO2 (all species in equilibrium) | 6.00E-05 | 1.00E-04 |
| for[e] | Formate | 3.26E-02 | 3.26E-02 |
| suc[e] | Succinate | 4.23E-03 | 4.23E-03 |
| nh3[e] | Ammonia | 5.50E-04 | 5.50E-02 |
| frc[e] | Fructose | 5.55E-03 | 5.55E-03 |
| pi[e] | Orthophosphate | 5.24E-02 | 5.24E-02 |

#### 2. MDF framework used to evaluate max-min Driving Force for each EFM

The framework for thermodynamic max-min driving force analysis was adapted from Noor et al., 2014. The linear programming problem was formulated as described below:

$$\mathbf{c} \equiv \begin{pmatrix} 0_{(n)} \\ 1 \\ 0_{(n)} \\ 0_{(n_{trnsp})} \end{pmatrix} \in \mathbb{R}^{(n+1+n+n_{trnsp})}$$

$$\mathbf{A} \equiv \begin{pmatrix} S^T_{no\_ex} & 1_{(m_{no_{ex}},1)} & S^T_{no\_ex} & S^T_{trnsp\_no\_ex} \\ I_{(n,n)} & 0_{(n,1)} & 0_{(n,n)} & 0_{(n,n_{trnsp})} \\ -I_{(n,n)} & 0_{(n,1)} & 0_{(n,n)} & 0_{(n,n_{trnsp})} \\ 0_{(n,n)} & 0_{(n,1)} & I_{(n,n)} & 0_{(n,n_{trnsp})} \\ 0_{(n,n)} & 0_{(n,1)} & -I_{(n,n)} & 0_{(n,n_{trnsp})} \\ 0_{(n_{trnsp},n)} & 0_{(n_{trnsp},1)} & 0_{(n_{trnsp},n)} & I_{(n_{trnsp},n_{trnsp})} \\ 0_{(n_{trnsp},n)} & 0_{(n_{trnsp},1)} & 0_{(n_{trnsp},n)} & -I_{(n_{trnsp},n_{trnsp})} \\ S^T_{ratios} & 0_{(n_{ratios},1)} & 0_{(n_{ratios},n)} & 0_{(n_{ratios},n_{trnsp})} \\ -S^T_{ratios} & 0_{(n_{ratios},1)} & 0_{(n_{ratios},n)} & 0_{(n_{ratios},n_{trnsp})} \end{pmatrix} \in \mathbb{R}^{(m_{no_{ex}}+n+n+n+n+n_{trnsp}+n_{trnsp}+n_{ratios}+n_{ratios})\times(n+1+n+n_{trnsp})}$$

$$\mathbf{x} \equiv \begin{pmatrix} \ln[m]_{(n)} \\ B \\ d_r G'_{(n)} \\ d_t G'_{(n_{trnsp})} \end{pmatrix} \in \mathbb{R}^{(n+1+n+n_{trnsp})}$$

$$\mathbf{b} \equiv \begin{pmatrix} 0_{(m_{no_{ex}},1)} \\ \ln[m]_{min_{(n)}} \\ -\ln[m]_{max_{(n)}} \\ d_f G'^0{}_{min}/RT_{(n)} \\ -d_f G'^0{}_{max}/RT_{(n)} \\ d_t G'^0/RT_{(n_{trnsp})} \\ -d_t G'^0/RT_{(n_{trnsp})} \\ \ln CoFR_{min_{(n_{ratios})}} \\ -\ln CoFR_{max_{(n_{ratios})}} \end{pmatrix} \in \mathbb{R}^{(m_{no_{ex}}+n+n+n+n+n_{trnsp}+n_{trnsp}+n_{ratios}+n_{ratios})}$$

With n being the number of metabolites, $n_{trnsp}$ the number of transport reactions, $n_{ratios}$ the number of cofactor ratios, $m_{no\_ex}$ the number of reactions excluding exchange reactions, the biomass reaction as well as the PHB synthesis reaction.

#### 3. Strain Construction

The plasmid pBBR1-MCS-2 was linearised with primers NC93 + NC94 and three plasmids were constructed via HiFi cloning (New England Biolabs). The fragments containing RFP,

AclAB or XFPK were amplified from the pCM271RFP, pCM271_aclAB and pCM271_xfp_pta plasmids (Nuha, 2019) using the primers NC29 + NC65. The resulting vectors were termed HSLP 001, HSLP 002 and HSLP 003 and were transformed into XL1-blue *E.coli* to make the stocking strains HSL147, HSL224 and HSL225. The three plasmids were subsequently transferred to SL-17 E.coli and conjugated into C. necator H16 to form the mutant strains HSL 002 (containing RFP), HSL 229 (containing AclAB), and HSL 230 (containing XFPK). Plasmids were confirmed via sequencing using the sequencing primers listed in the primer table. Screening primers NC107 + NC108/109/110 were used to confirm the plasmids presence in the *C. necator* strains along with measurements of RFP fluorescence, aclAB enzyme activity and western blot analysis respectively for HSLP 001, HSLP 002, HSLP 003.

#### Supplementary Figures

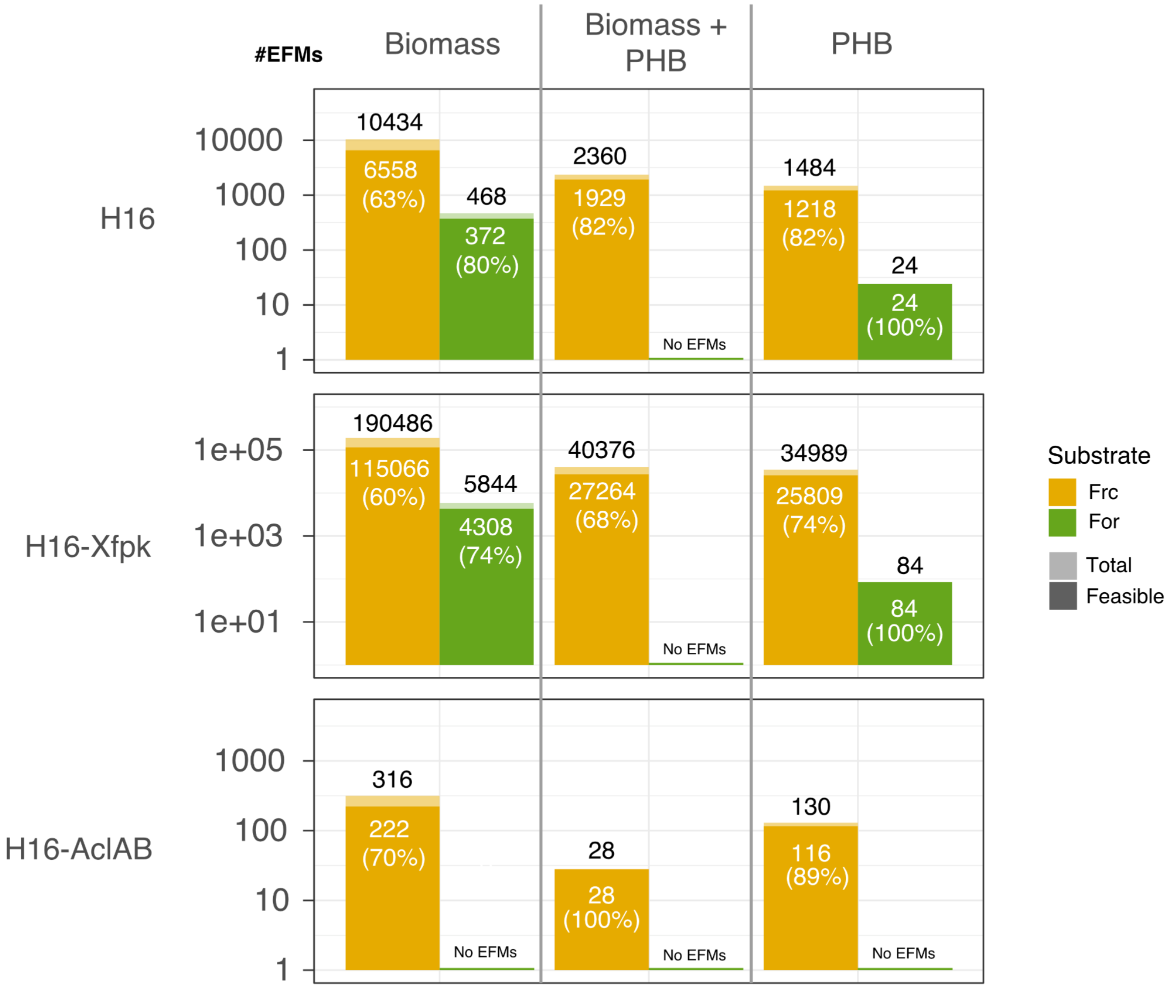


**Figure S1: Flexibility of *C. necator*'s core metabolism.** Total number of EFMs (lighter shade, black number) and number of thermodynamically feasible EFMs (darker shade, MDF > 0, white number) creating biomass, PHB or both, for each strain for fructose and formate. Percentage numbers indicate feasible fraction of total EFM number. The EFM numbers for the mutant strains Xfpk and AclAB represent EFMs with non-zero flux through the corresponding reaction(s). Percentage of feasible EFMs is written in the bars.

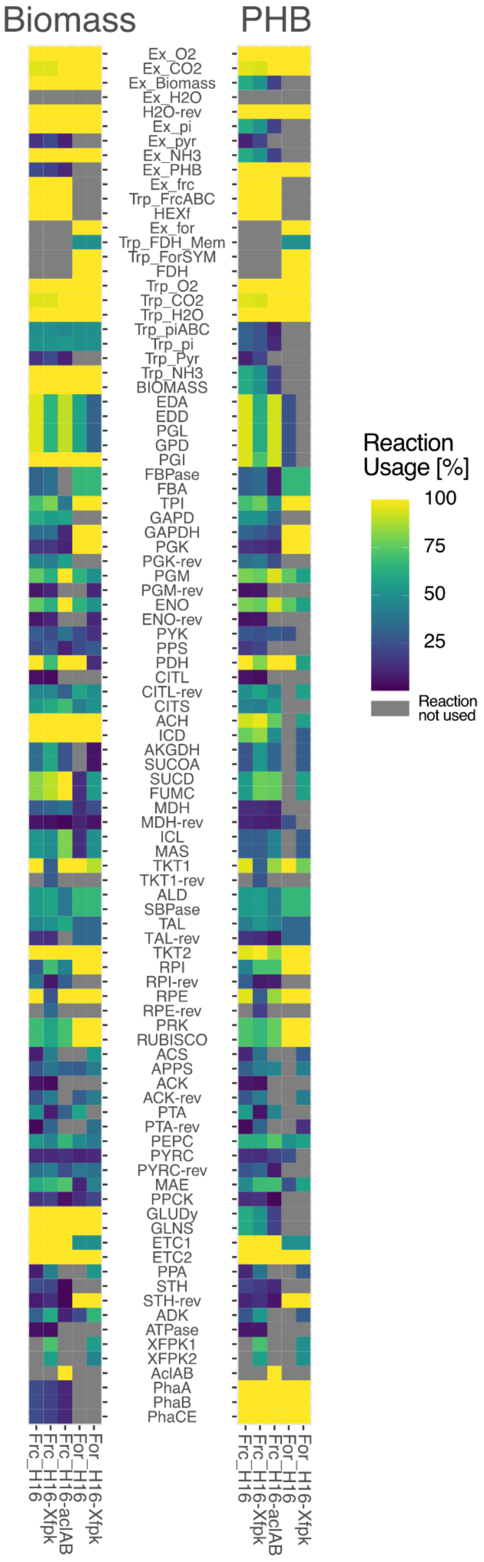


**Figure S2: Reaction usage of *C. necator’s* core metabolism in dependency of the strain and substrate.** Reaction usage as percent of total number of thermodynamically feasible EFMs for biomass and PHB production. A reaction usage of 100% signifies essentiality.

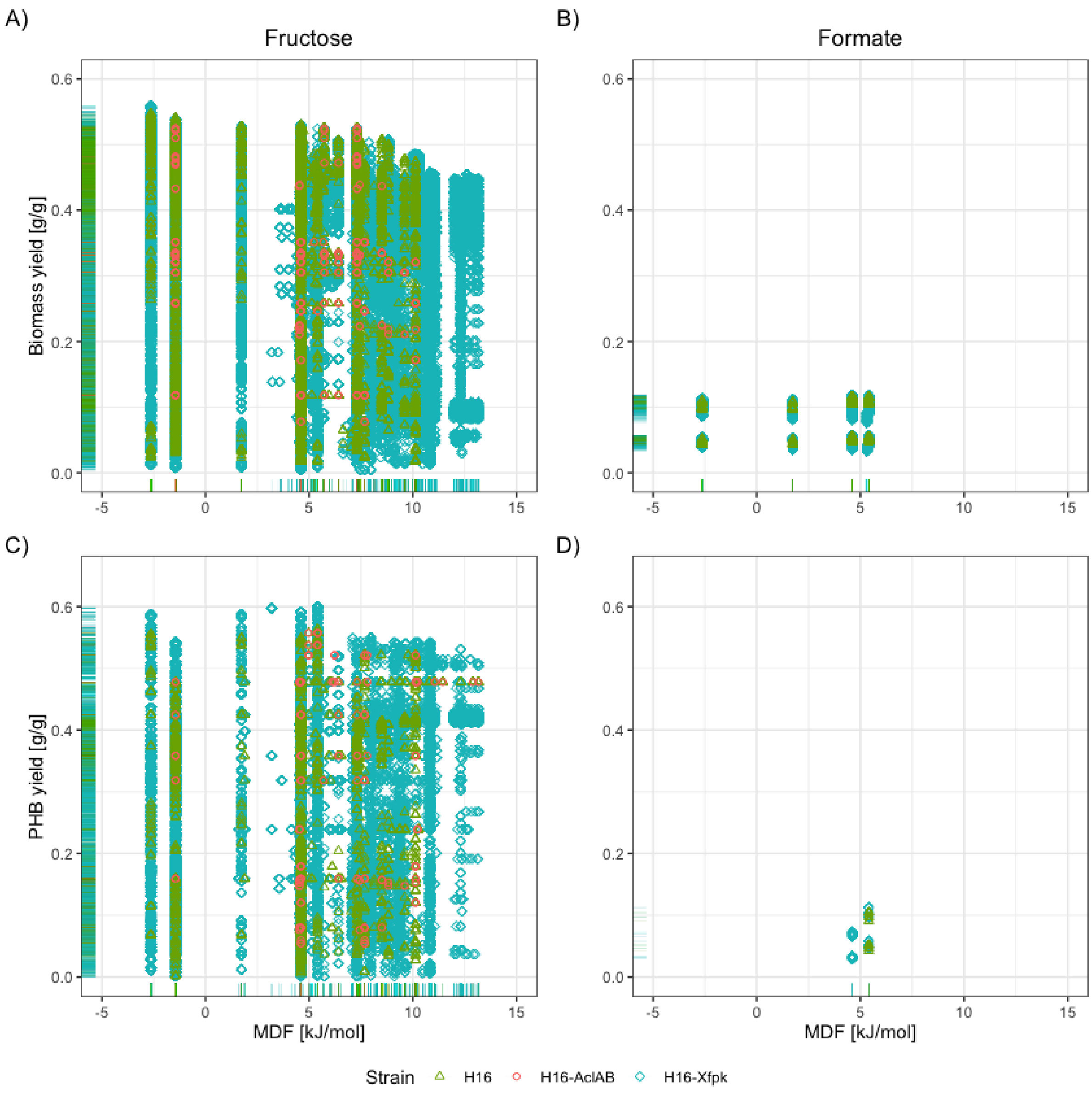


**Figure S3: Relationship of Yields and MDFs for biomass and PHB producing EFMs.** For each EFM**,** the biomass (A-B) and PHB (C-D) yield on the different substrates (g/g substrate) is plotted over the MDF. Only EFMs with non-zero biomass of PHB yields are shown in the subplots, respectively. Rug-plots on the bottom and left side of each subplot show the distribution of the MDF and yield values individually.

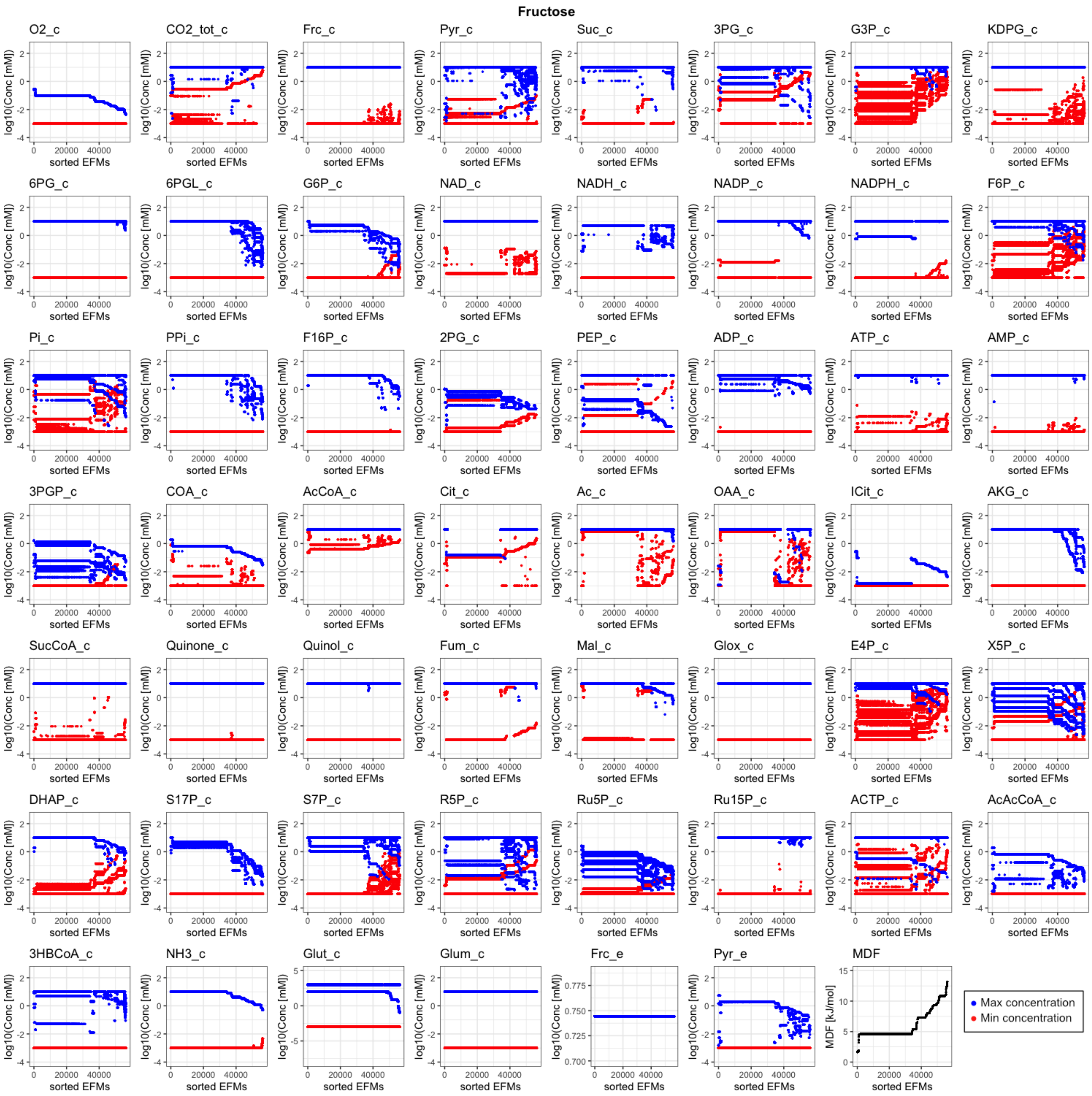


**Figure S4: Metabolite concentration ranges for EFMs producing PHB under fructose utilization.** Concentration ranges for each metabolite for each EFM utilizing fructose for PHB production, resulting from Energy Variability Analysis. EFMs, and thereby ranges, are sorted after the corresponding MDF. EFMs with the same MDF are sorted by the mean DF over all reactions in the corresponding EFM. Many metabolite concentrations become more constrained with increasing MDF. EFMs with no data point indicate that metabolite is not used.

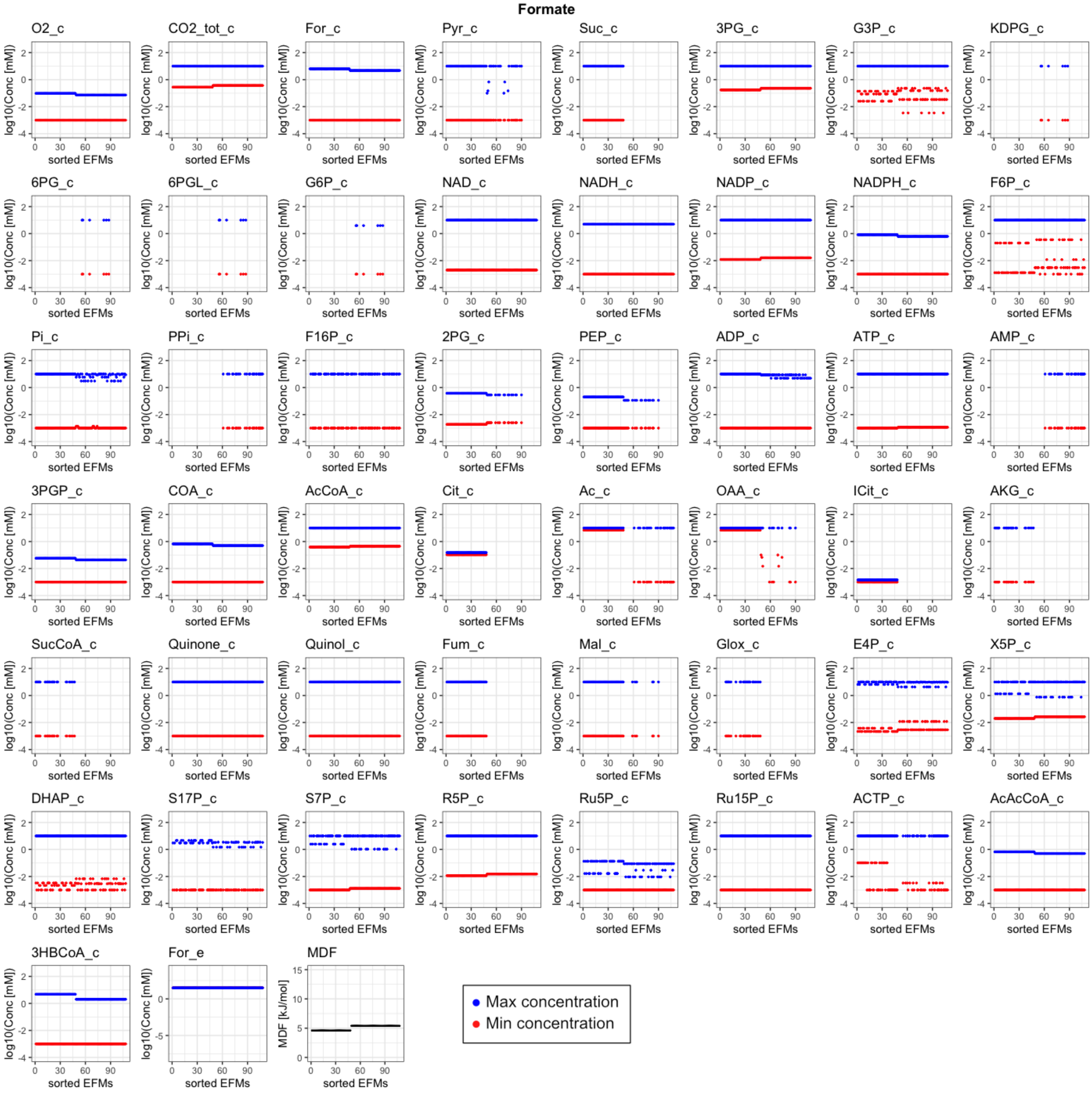


**Figure S5: Metabolite concentration ranges for EFMs producing PHB under formate utilization.** Concentration ranges for each metabolite for each EFM utilizing formate for PHB production, resulting from Energy Variability Analysis. EFMs, and thereby ranges, are sorted after the corresponding MDF. EFMs with the same MDF are sorted by the mean DF over all reactions in the corresponding EFM. EFMs with no data point indicate that metabolite is not used.

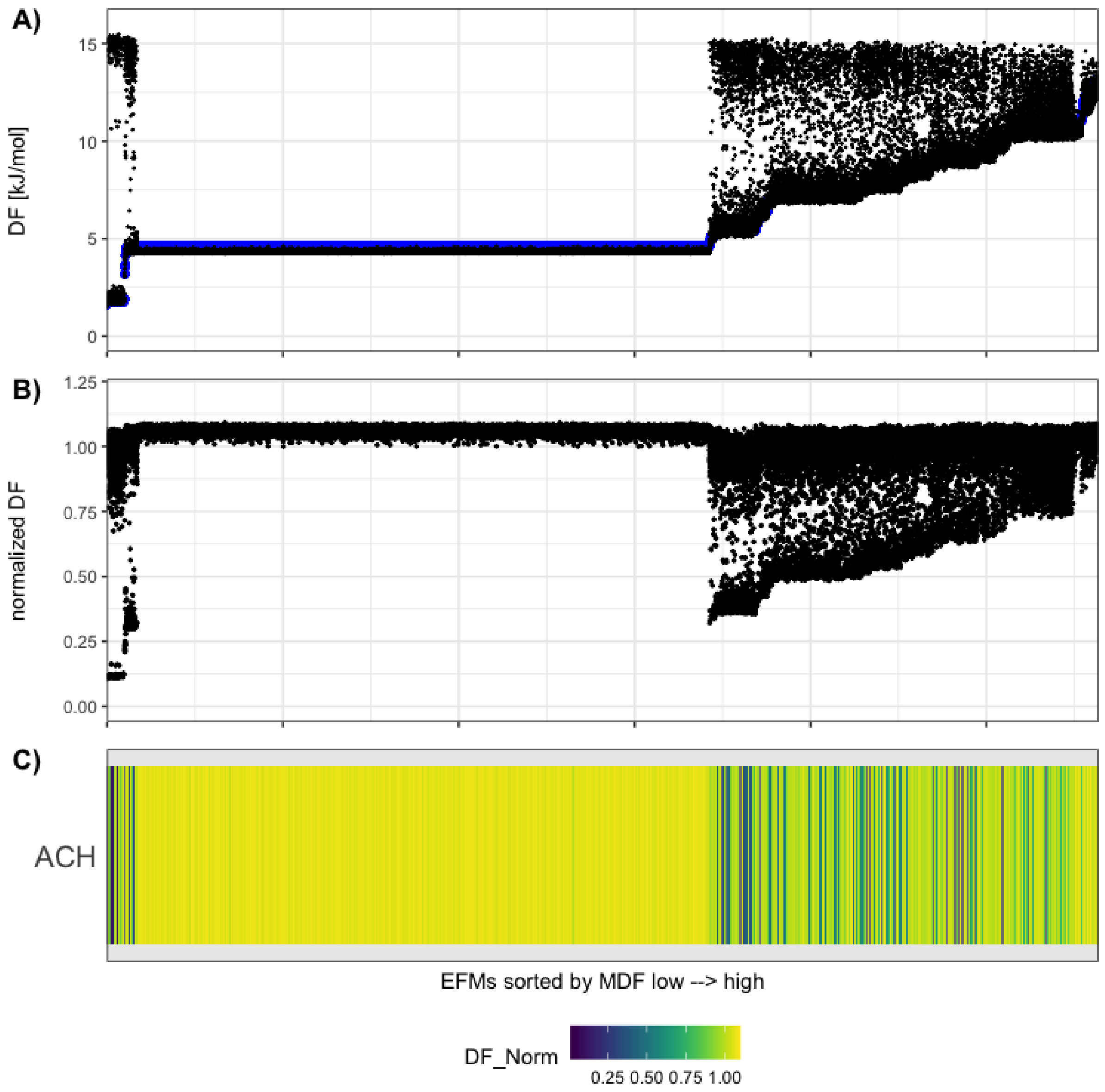


**Figure S6: Example for derivation of the DF_Norm value.** Derivation DF_Norm value as used in heatmaps for Figure 4 A and Supplementary Figures S7-S9. **A)** Median DF for ACH for each EFM analyzed for PHB production on fructose, derived from random sampling. Also displayed is the MDF for each EFM (blue line). **B)** DF_Norm calculated by dividing the MDF value for each EFM by the median DF for ACH for each EFM. A value of 1 represents DF = MDF. **C)** Translation of B) into a heatmap. Gray colors indicate that ACH was not used in that specific EFM.

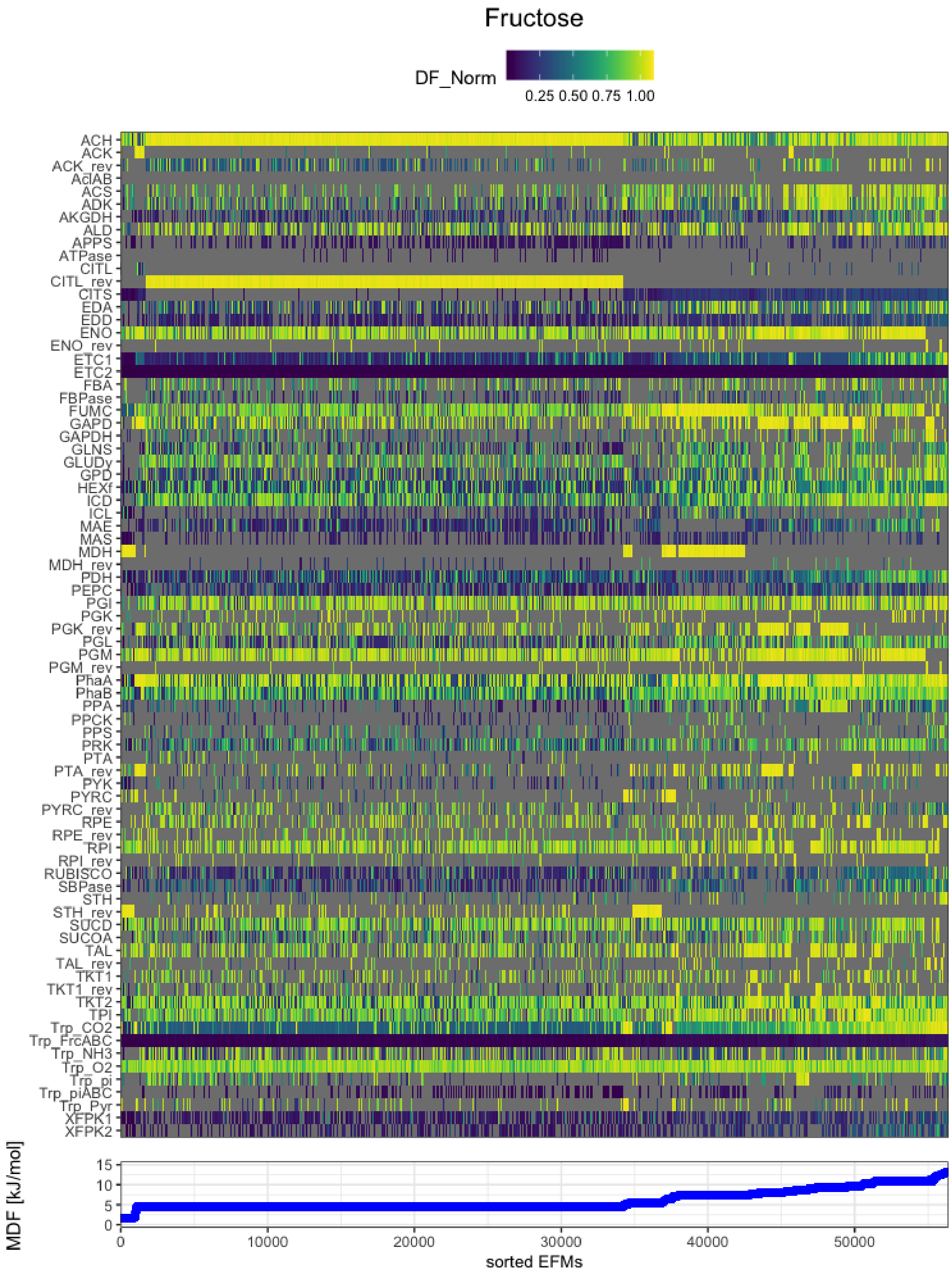


**Figure S7: DF_Norm for all reactions for all EFMs producing PHB under fructose utilization.** Reaction driving forces for each reaction for each EFM utilizing fructose, normalized with the corresponding EFM's MDF. The grey color indicates reactions not being used in the respective EFM.

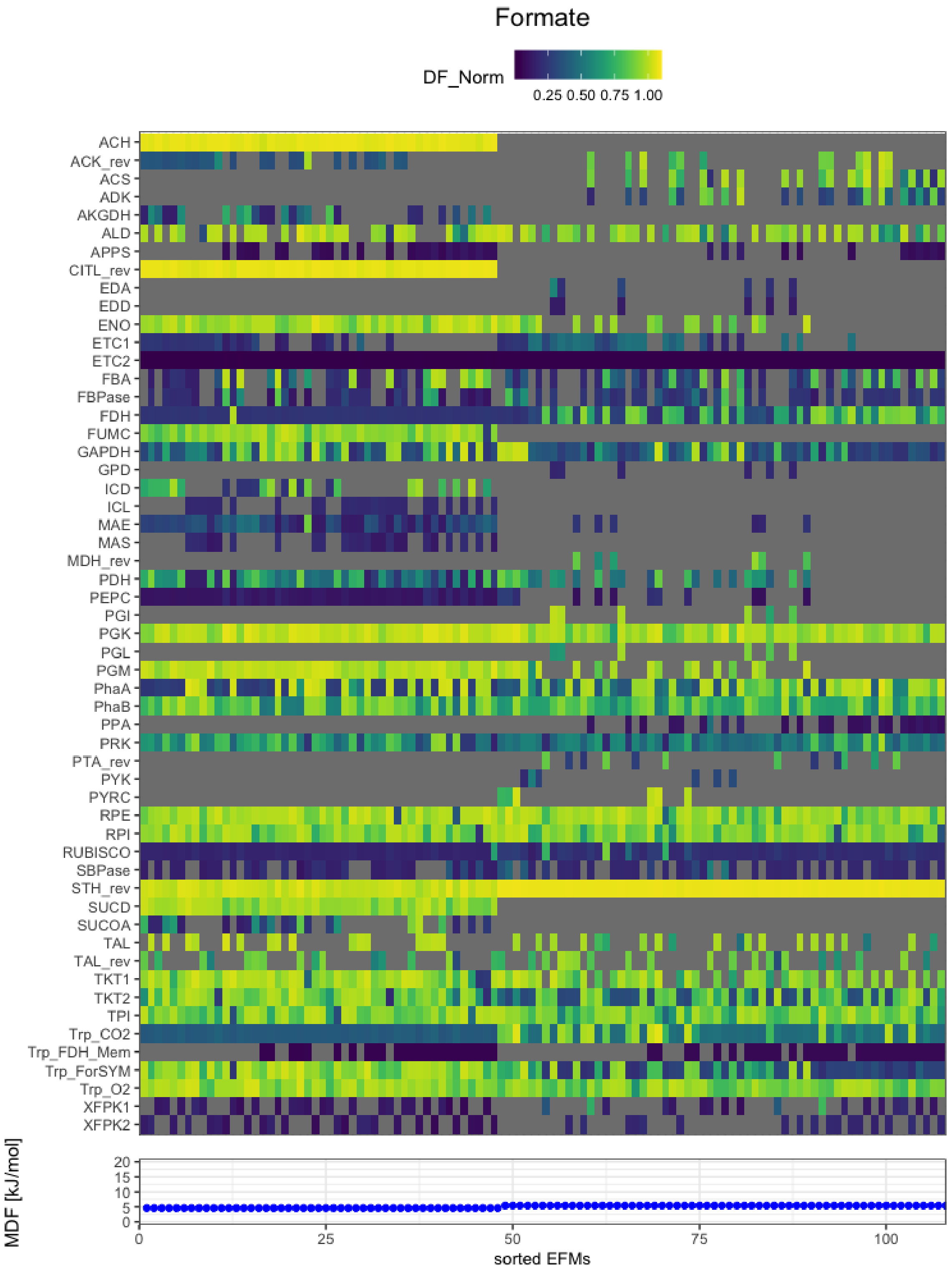


**Figure S8: DF_Norm for all reactions for all EFMs producing PHB under formate utilization.** Reaction driving forces for each reaction for each EFM utilizing formate, normalized with the corresponding EFM's MDF. The gray color indicates reactions not being used in the respective EFM.

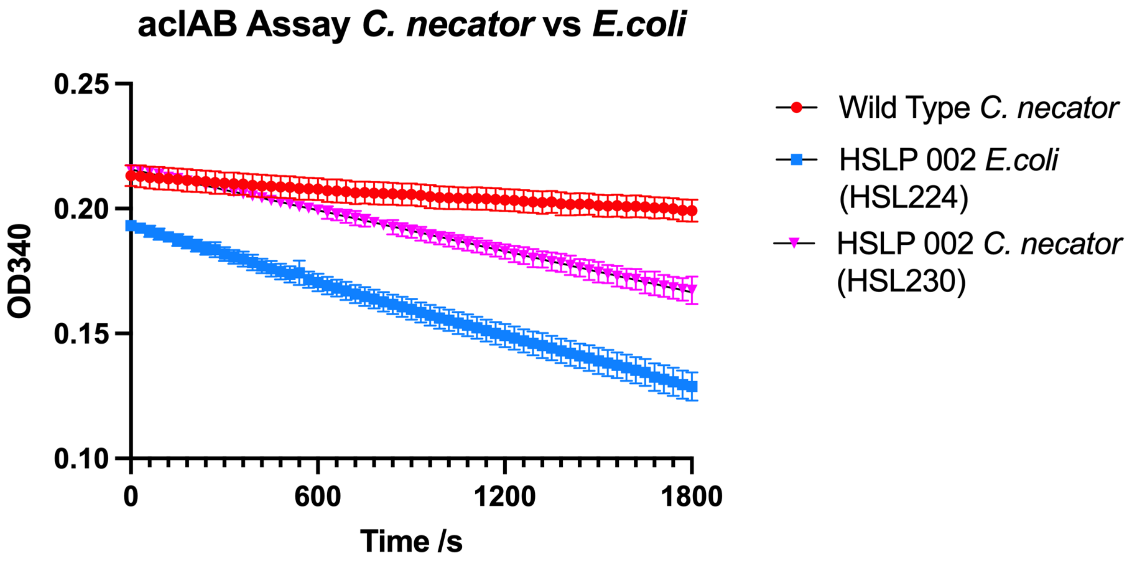


**Figure S9: Enzymatic assay of AclAB activity from extracts of recombinant *C. necator* and *E. coli*.** *C. necator* or *E. coli* strains containing the HSLP 002 plasmid were grown in shake flask cultures (LB) and induced with 1 mM arabinose. After overnight incubation (OD600 =1), 10 mL of cell culture was harvested, lysed, and soluble protein extracted. Soluble protein (5 ug) was assayed for AclAB activity using a TECAN plate reader (Methods).

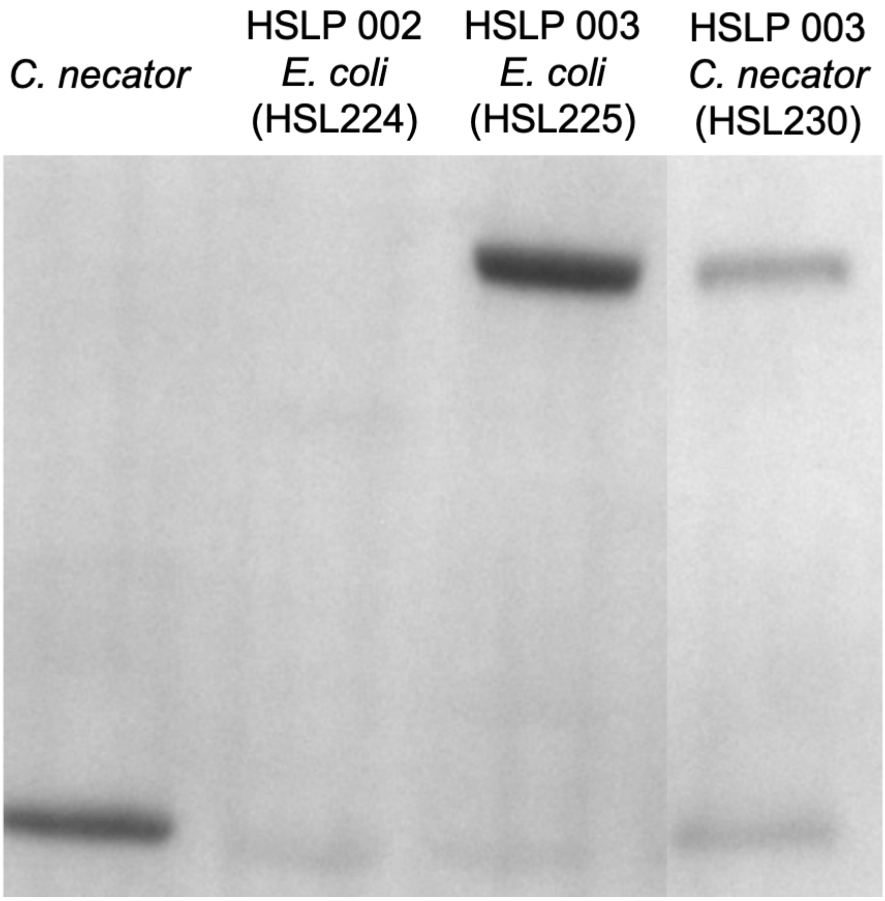


**Figure S10: Western blot of Xfpk expression in recombinant *C. necator* and recombinant *E. coli*.** *C. necator* or *E. coli* strains were grown in shake flask cultures (LB) and induced with 1 mM arabinose. After overnight incubation (OD600 =1), 10 mL of cell culture was harvested, lysed, and soluble protein extracted. Soluble protein (10 ug) was loaded for Western blotting using an anti 6-His primary antibody (Methods). A clear band was observed from *C. necator* and *E. coli* strains containing the Xfpk-encoding HSLP 003 plasmid in both *E.coli* and *C. necator*. Wild type *C. necator* or *E. coli* expressing AclAB (strain HSL224) were used as negative controls and did not show a Xfpk band. The band arising from wild type *C. necator* extracts was reproducible but the protein was not identified.

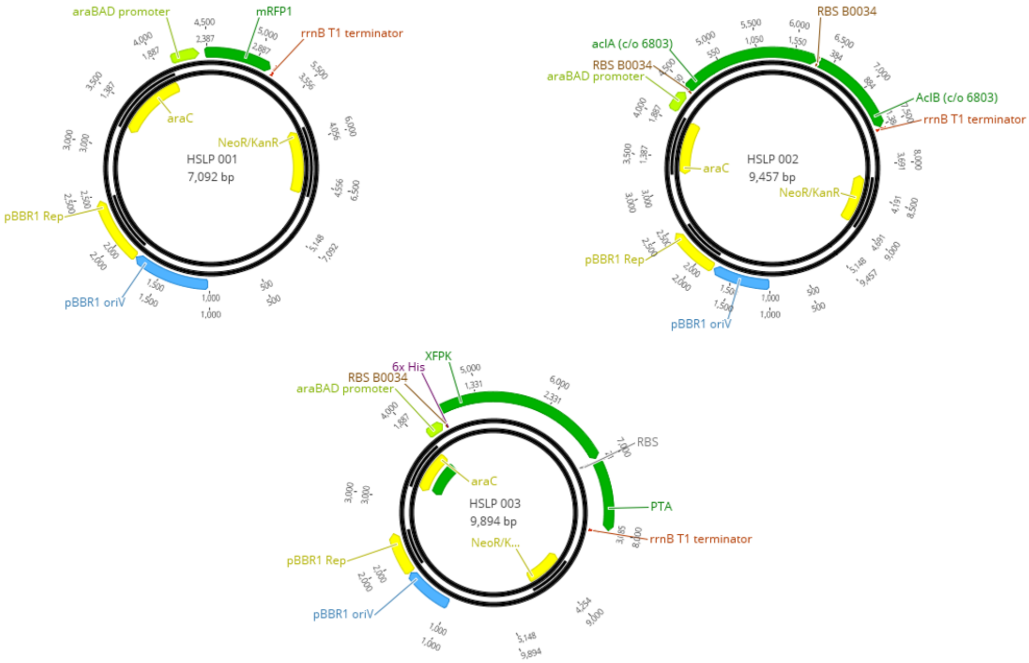


**Fig S11: Plasmid maps**

Plasmid maps of the three arabinose inducible plasmids used in this study. HSLP 001 is the Rfp, HSLP 002 contains the AclAB and HSLP 003 contains the Xfpk and PTA genes.
